# Astrocyte-neuron mitochondrial transfer via mitoEVs supports neuronal energy metabolism and is impaired in early Alzheimer’s disease

**DOI:** 10.64898/2026.03.09.710050

**Authors:** Liesbeth Voorbraeck, Jesus Alarcon-Gil, Romain Giraud, Francesca Pozzobon, Maria João Pereira, Shubin Guo, Zhibai Cao, Krizia Distefano, Dara K. Mohammad, Oscar P. B. Wiklander, Mite Mijalkov, Joana B. Pereira, Doste R. Mamand, Maria Ankarcrona, Luana Naia

## Abstract

**Background:** Mitochondrial dysfunction is an early and central feature of Alzheimer’s disease (AD). In particular, intercellular mitochondrial transfer has emerged as a mechanism of neuronal support in brain injury and neurodegeneration. However, pathways governing astrocyte-to-neuron transfer and its role in AD pathogenesis remain unknown.

**Methods:** Using the *App^NL-G-F^* knock-in AD model, we combined high-resolution 4D live-cell imaging with quantitative fluorescence-based reporters to assess synaptic function and mitochondrial network dynamics in neurons and astrocytes. Direct and extracellular vesicle (EV)-restricted neuron-astrocyte co-culture systems were used to investigate bidirectional mitochondrial transfer. We performed the first in-depth structural, proteomic, and functional characterization of astrocyte-derived mitochondrial extracellular vesicles (mitoEVs) using cryo-electron microscopy, quantitative mass spectrometry, and bioenergetic analyses to define their cargo composition and metabolic effects.

**Results:** We identified cell-type–specific mitochondrial remodeling in early AD, with compartmentalized synaptic energy deficits in neurons and hyperdynamic, less interconnected, yet metabolically preserved networks in astrocytes, preceding global bioenergetic decline. Bidirectional mitochondrial transfer between astrocytes and neurons, also at axonal terminals, was mediated by specialized mitoEVs but significantly reduced in the *App^NL-G-F^*model. Comprehensive proteomic and functional profiling revealed that WT astrocyte-derived mitoEVs are enriched in inner membrane and matrix proteins, supporting oxidative phosphorylation, lipid and amino acid metabolism, and redox homeostasis. In contrast, *App^NL-G-F^* mitoEVs are selectively depleted of respiratory and fatty acid oxidation components and exhibit impaired respiration with reduced Complex IV activity. Functionally, WT mitoEVs promote mobilization of abnormal accumulation of lipid droplets in *App^NL-G-F^*neurons, restore fatty acid oxidation, and increase neuronal bioenergetics, including at the synapses. In contrast, disease-derived mitoEVs fail to engage these pathways.

**Conclusions:** Together, these findings identify mitoEV-mediated mitochondrial transfer as a glia-to-neuron metabolic pathway compromised in early AD and reveal a coordinated role for oxidative phosphorylation and fatty acid oxidation in supporting synaptic energy homeostasis.

## INTRODUCTION

Alzheimer’s disease (AD), the leading cause of dementia worldwide, is a progressive and multifactorial neurodegenerative disorder with profound social and economic consequences. It is characterized by the accumulation of extracellular amyloid β-peptide (Aβ) aggregates, intracellular neurofibrillary tangles formed by hyperphosphorylated tau, extensive neuroinflammation, and selective neuronal loss. Beyond these alterations, increasing evidence places mitochondrial abnormalities among the earliest events in AD, suggesting that they should be considered in the etiology and treatment of the disease [1].

Mitochondria are double-membraned, dynamic, and signaling organelles optimized for supplying energy in the form of ATP. Previously, our laboratory has shown that Aβ directly interacts with and is internalized into mitochondria via the translocase of the outer membrane (TOM) [2], contributing to bioenergetic impairments [3]. Transcriptomic and functional studies in pre-symptomatic *App* knock-in mouse models indicate significant alterations in oxidative phosphorylation (OXPHOS; the final biochemical pathway in ATP production) associated with increased production of reactive oxygen species (ROS) and Ca^2+^ mishandling [4]. These mitochondrial alterations occur concurrently with the loss of high-frequency gamma oscillations in the hippocampus, which are important for memory formation and depend on robust mitochondrial function [5, 6].

Mitochondrial functions are tightly bound to mitochondrial network dynamics – especially in highly polarized cells such as neurons – including trafficking, fusion and fission, cristae remodeling, and generation of mitochondrial-derived vesicles (MDV) [7]. Recent reports have shown that mitochondria can operate beyond the boundaries of a single cell, undergoing dynamic physical transfer from cell to cell. Mitochondrial transfer from neighboring cells has been shown under a variety of circumstances – to promote immune evasion by cancer cells through mitochondria hijacking [8, 9], to foster metabolic adaptation to lipotoxic stress elicited by obesity [10], and to protect ischemic neurons after stroke [11].

Within the central nervous system, however, data on mitochondrial transfer are scarce and poorly defined. Astrocytes have emerged as probable mitochondrial donors, given their proximity to synapses and their acknowledged role in neuronal metabolic coupling [11, 12]. While previous studies propose that horizontal mitochondrial transfer from astrocytes to neurons can enhance metabolic activity and contribute to the upregulation of cell survival signals after stroke [9, 11], reactive microglia and astrocytes also release dysfunctional fragmented mitochondria and propagate neurodegeneration [13].

The molecular mechanisms regulating mitochondrial transfer remain incompletely understood. The best described routes involve either tunneling nanotubes (TNTs) [12] or extracellular vesicles (EVs) [14]. EVs vary in size, cargo, and origin, but ground-breaking studies from Efrat Levy’s lab led to the identification of a specific subpopulation of double-membraned EVs enriched in mitochondrial proteins, with intact mitochondrial membrane potential and lacking tetraspanin proteins commonly found on the membrane of exosomes, which were named mitovesicles (hereafter, mitoEVs) [15, 16]. MitoEVs appear to originate from mitochondrial membrane protrusions that undergo scission, a process regulated by retromer proteins (VPS26/29/35), Miro-GTPases, and fission proteins such as dynamin-related Protein 1 (Drp1) and mitochondrial fission factor [17]. In fact, mitoEVs are likely MDVs that, instead of being delivered to lysosomes or peroxisomes for mitochondrial quality control (their first recognized role), are targeted to multivesicular bodies for release into the extracellular space [17]. MitoEVs have been described to carry membrane transporters for metabolites, they appear to contain a functional ATP synthase and retain their electrochemical potential [18], as well as elicit antioxidant responses in other cells and tissues [10, 19]. While these features support the idea that mitoEVs, when transferred, can promote protection in host cells, specific cargo may also contribute to toxicity. Both Aβ and tau have been found in EVs [20] and are likely involved in the spread of pathology, including early synaptic impairment [21]. Additionally, the surface of mitoEVs seems to be enriched in monoamine oxidase type B (MAO-B), a marker of reactive astrocytes, which can affect synaptic plasticity when delivered to brain slices [22]. Therefore, mitoEVs may represent a novel class of context-dependent intercellular signals capable of either supporting or disrupting neuronal homeostasis and thus require further investigation.

In this study, we ask whether defective astrocyte–neuron mitochondrial transfer contributes to the synaptic bioenergetic decline observed in the *App^NL-G-F^* knock-in model of AD. We identify mitoEVs as mediators of astrocyte-to-neuron transfer, including at axonal terminals, and show that this process is reduced under AD conditions. Proteomic and respirometry profiling reveal genotype-specific astrocytic mitoEV signatures, with decreased enrichment of inner mitochondrial membrane and matrix proteins in *App^NL-G-F^* mitoEVs, associated with reduced lipid metabolism and OXPHOS capacity. In recipient neurons, WT mitoEV cargo promotes fatty acid utilization, leading to reduced lipid droplet accumulation, and restores synaptic ATP levels, in a process dependent on mitochondrial fatty acid entry. In contrast, *App^NL-G-F^*-derived mitoEVs fail to rescue neuronal metabolic competence. Together, these findings identify mitoEV-mediated mitochondrial transfer as a key pathway of astrocyte-to-neuron metabolic support that is compromised in early AD.

## METHODS AND MATERIALS

### Animal and ethical considerations

*App* knock-in mice harboring Swedish (K670/M671L), Beyreuther/Iberian (I716F) and Arctic (E693G) mutations [23] (hereafter called *App^NL-G-F^*) and wild-type (WT) mice derived from heterozygous *App^NL-G-F^* breeding, all with C57B6/J genetic background, were used. Animals were bred and housed at the Comparative Medicine’s Annex (KM-A) at Karolinska Institutet under conditions of controlled temperature (22–23°C) and 12:12 h light/dark cycles and specific pathogen-free conditions. Cages were enriched in corn-husk nesting material and paper rolls. Food and water were available *ad libitum*. All experimental procedures were carried out in accordance with the guidelines of the Institutional Animal Care and Use Committee and the European Community directive (2010/63/EU), and with protocols approved by the Regional Ethics Review Board (authorizations no. 12779/2019 and 14573-2024), in line with the ARRIVE guidelines. All efforts were made to minimize animal suffering and to reduce the number of animals used. For *ex vivo* experiments, four 3-month-old female mice per genotype were used and were perfused under anesthesia. For primary cultures, females of reproductive age (2–6 months old) were used for breeding. Mating occurred overnight (∼15 h) with two females paired with one male. Mice were euthanized by cervical dislocation under anesthesia (5% isoflurane). Brains were collected and maintained in Hibernate-E medium (Thermo Fisher Scientific) until dissection.

### Primary cell culture, viral transduction and transfection

Primary cortico-hippocampal neurons and astrocytes were collected from WT and *App^NL-G-F^* mouse embryos (from both sexes) at embryonic day 17. Dissection was performed in ice-cold Hank’s balanced salt solution (HBSS, Thermo Fisher). Neuronal cultures were obtained by trypsinizing the tissue in 0.06% trypsin/EDTA (prepared in HBSS/BSA) at 37°C for 13 minutes. Trypsinization was stopped by adding 10% heat inactivated (HI) fetal bovine serum (FBS, Thermo Fisher). Tissues were mechanically dissociated in 0.2 mg/ml DNase I (prepared in HBSS/BSA) using a sterile flame-polished Pasteur pipette. Neurons were cultured in Poly-D-lysine-coated plates in neurobasal medium supplemented with 2% B-27 (Thermo Fisher) and 1% glutaMAX (Thermo Fisher). To obtain astrocyte cultures, two embryonic cortices/hippocampus were mechanically disassociated in 10 mL of DMEM/F12 medium (Thermo Fisher) supplemented with 10% HI FBS, 1% N2 (Thermo Fisher), then filtered with a 40 μm cell strainer and transferred to a non-coated T75 flask. Cell media were changed at DIV (day *in vitro*) 3 and DIV7. At DIV 7, 0.5 μM of PLX-3397 (MedChemExpress), a CSF1R inhibitor, was added to the culture to deplete microglia. At DIV13, when cells reached confluence, astrocytes were passaged. Briefly, astrocytes were dissociated with 0.25% Trypsin/EDTA. Trypsin was blocked with FBS-supplemented DMEM/F12, and astrocytes were collected after centrifugation at 500 *xg* for 5 min. Astrocytes were counted and then seeded in plates at the desired density. Both astrocytes and neurons were incubated at 37 °C and 5% CO_2_/95% air for further experiments.

Neurons were transfected or transduced at DIV4, whereas astrocytes were transfected or transduced after the first passage. The following lentiviral (LV) and adeno-associated viral (AAV) vectors were used: LV-mitoDsRed, produced at the Karolinska VirusCore, and AAV1-GFAP-mitoDsRed produced by Vector Biolabs using the pLV-mitoDsRed vector (a kind gift from Pantelis Tsoulfas; #44386, Addgene; http://n2t.net/addgene:44386; RRID: Addgene_44386) and applied at a multiplicity of infection (MOI) of 10 (for LV) and at a dilution of 1:100 from a titer of 1 × 10^12^ GC/mL (for AAV); AAV1-eSYN-mEmerald-Mito-7-WPRE, produced by Vector Biolabs using the mEmerald-Mito-7 plasmid (a gift from Michael Davidson; #54160, Addgene; http://n2t.net/addgene:54160; RRID: Addgene_54160) and used at a dilution of 1:500 from a titer of 1 × 10⁹ GC/mL; AAV2-eSYN-GFP-WPRE and AAV2-CAG-CFP, produced by Vector Biolabs and used at a dilution of 1:100 from a titer of 1 × 10^13^ GC/ml and 1 × 10⁹ GC/mL, respectively.

Transfections were performed using Lipofectamine 2000 (Thermo Fisher Scientific) according to the manufacturer’s instructions. The following plasmids were used: mKeima-Red-Mito-7 (a gift from Michael Davidson; #56018, Addgene; http://n2t.net/addgene:56018; RRID: Addgene_56018); CMV::SypHy A4 (a kind gift from Prof. Leon Lagnado, University of Sussex; #24478, Addgene; http://n2t.net/addgene:24478; RRID: Addgene_24478); and sypI-ATeam1.03 (YFP/CFP) and the sypI-ATeam1.03 R122K/R126K mutant, kindly provided by Daniel Gitler (Zlotowski Center for Neuroscience, Israel).

<u>Co-cultures in microfluid chambers</u>: silicone microfluidic chambers (Xona Microfluidics) were mounted on squared 22x22 mm poly-D-coated (0.5 mg/mL) coverslips. 1x10^5^ neurons in 50 µl of media were plated in one compartment near the microgrooves and cultured for 5 days, followed by transduction with the AAV1-eSYN-mEmerald-Mito-7-WPRE AAVs for 48h. At DIV6, 3.2x10^5^ mitoDsRed-transduced astrocytes were plated homogeneously in the other compartment, and co-cultures were maintained for 48 h.

<u>Co-cultures in Boyden chambers</u>: To study astrocyte-to-neuron mitoEVs transfer, primary neurons were plated on a coverslip and infected with the AAV1-eSYN-mEmerald-Mito-7-WPRE AAVs for 48h. MitoDsRed-transduced astrocytes were plated in transwell Boyden chambers with a membrane of 0.4 µm pore (Falcon) and placed above neurons at DIV10. After 48 hours of co-culturing, neurons were labelled with 20 µM CellTracker Blue (Thermo Fisher Scientific) at 37°C for 45 minutes. To study neuron-to-astrocyte mitoEVs transfer, neurons were transduced with mitoDsRed. After 48 h of co-culturing, performed as described above, the mitochondrial network in astrocytes was labeled with TOM20.

### Mitochondrial movement in axons

MitoDsRed-transfected neurons were washed and incubated in artificial cerebrospinal fluid (ACSF; NaCl 124 mM, KCl 2.5 mM, NaH_2_PO_4_ 1.25 mM, glucose 10 mM) supplemented with 1.5 mM CaCl_2_ and 1.5 mM MgCl_2_, and mitochondrial movement studies were carried out at 37°C. Live imaging was performed using a 63x objective with NA=1.4 in the LSM980-Airy confocal microscope (Zeiss). Acquisitions were performed with a 150 µm pinhole aperture, and images were captured every 5 s for a total of 120 frames. Histograms were matched to the first frame to correct for fluorescence variations using the Bleach Correction plugin developed by Miura and Rietdorf, and time-lapse-dependent x-y drift was corrected using the TurboReg plugin. Mitochondrial movement analysis was done using the Kymograph Macro in Fiji (Rietdorf and Seitz, 2004). Regions of interest (ROIs) were designated using a segmented line that followed the mitochondrial trajectory across projections. Kymographs generated in a *x*-*y* dimension (distance vs time) were used to obtain the slope from which mitochondrial velocity was calculated.

### Mitochondrial 4D network imaging analysis in astrocytes

MitoDsRed-transduced astrocytes were imaged in DMEM media without phenol red at 37 °C with a Nikon CrEST X Light V3 Spinning Disk microscope with a 60X objective. Z-stack volumes with a 0.2 µm step size were acquired at 0.5 frames per s for 4 min. The original mono-channel time-lapse images were transferred to an Ubuntu Linux distribution, where we developed a pipeline in Jupyter Notebook (Anaconda, Python) to preprocess, deconvolute, segment, and analyze the time-lapse images. The analysis code is available in a GitHub repository and will be made publicly available upon publication. Detailed preprocessing parameters and computational steps are provided in the Supplementary Files.

### Mitochondrial membrane potential and cellular ATP levels

Neurons were incubated with 10 nM TMRM (Thermo Fisher) (non-quenching conditions) in ACSF with 26 mM NaHCO_3_ for 30 min at 37 °C. Under these conditions, TMRM retention by mitochondria was measured to estimate changes in mitochondrial membrane potential (Δψ*_m_*). After washing, neurons were imaged at 37 °C, and z-stack images were acquired using a 63x objective with NA=1.4 on the LSM980-Airy confocal microscope (Zeiss). The Fiji software was used to process the z-stack images into a 3D reconstruction and fluorescence intensity was measured.

Cellular ATP levels were measured using the CellTiter-Glo® Luminescent Cell Viability Assay (Promega). This assay triggers cell lysis and uses luciferase to catalyze a reaction that generates a luminescent signal that directly correlates with the ATP concentration in the sample. At DIV12, media from neurons plated in 96-well plates was removed and replaced by 100 µL of unbuffered DMEM 5030 media (Sigma Aldrich) supplemented with 10 mM sodium pyruvate (Thermo Fisher) and 2 mM L-Glutamine (pH 7.2-7.4), to enhance OXPHOS activity. Cells were incubated at 37 °C for 1 h. After incubation, 100 µL of CellTiter-Glo® reagent was added to the cells, which were then shaken for 2 min. Luminescence levels were measured in a plate reader and results were normalized to protein levels.

### RNA extraction and real-time PCR (qPCR)

RNA was extracted from astrocytes using the RNeasy® Mini Kit (Qiagen) by following the manufacturer’s instructions. RNA concentration was determined using the Nanodrop 2000 spectrophotometer (Thermo Scientific) and the RNA integrity was confirmed by A260/280 > 1.9. RNA was diluted to a final concentration of 10 ng/µL and converted into cDNA using the High-Capacity cDNA Reverse Transcription Kit (Applied Biosystems). The reaction was carried out in the S1000 thermocycler (Bio-Rad) with the following steps: 10 min at 25°C, 120 min at 37°C and 5 min at 85°C. qPCR was performed using the TaqMan™ Fast Advanced Master Mix (Applied Biosystems) with the primers listed in Supplementary Table S1 and run on the Applied Biosystems™ 7500 Fast qPCR System. Tubb3 was used as the housekeeping gene. Results are expressed as 2^ minus double delta Ct values (2^-ΔΔCt^).

### Synaptic ATP levels

Neurons transfected with the synaptic targeted FRET-based ATP sensor SypI-ATeam1.03 or with the ATP-insensitive mutant SypI-ATeam1.03^R122K/R126K^ were washed and imaged in ACSF with NaHCO_3_, 1.5 mM CaCl_2_ and 1.5 mM MgCl_2_. When indicated, neurons were treated with 4 μM etomoxir for 2 hours (Sigma Aldrich). The experiment was carried out at 37 °C, and Z-stack images were acquired with a Plan-Apochromat/1.4 NA 63x lens. Donor (mseCFP) was excited at 440 nm and its emission was measured at 460–500 nm before and after acceptor photobleaching; excitation to the acceptor (cpmVen) was performed at 514 nm, and emission was measured at 530–600 nm. Regional photobleaching was performed with the 514 nm laser.

3D reconstructions were performed using Fiji software. FRET ratio (F_cpmVen_/F_mseCFP_) for each region of interest was measured as described previously [24].

### Measurement of synaptic vesicle release with SypHy

SypHy is an exocytosis/endocytosis reporter localizing at synaptic vesicles, which uses the superecliptic pHluorin fused to the second intravesicular loop of synaptophysin. SypHy-transfected neurons were washed with ACSF and imaging was performed at 37 °C using a 63x objective with NA=1.4 in the LSM980-Airy confocal microscope (Zeiss). Images were acquired at 10 s intervals, and after 1 min of basal recording, KCl (50 mM) was added to the medium to induce depolarization and vesicle release. The fluorescence intensity of each ROI across the various frames was normalized to its pre-stimulation intensity and plotted over time. Radiometric analysis of final fluorescence to initial fluorescence (f/f0) was calculated by dividing the final fluorescence value of the ROI by the initial ROI fluorescence for each curve analyzed.

### Isolation of synaptosome fractions

Primary neurons were plated as described before and a 6-well tissue plate was used per condition at a seeding density of 500,000 cells/well. Isolation of synaptosome fractions and enrichment of post-synaptic densities (PSDs) was performed according to a published protocol[25]. First, cells are lysed in Buffer A (0.32 M sucrose, 4 mM HEPES-NaOH, pH 7.4) and homogenized. The homogenate was subjected to 1,000 × g for 10 min (4 °C) to remove nuclei and large debris. The supernatant, containing crude synaptosomes, was further centrifuged at 10,000 × g for 15 min (4 °C), yielding a pellet enriched in synaptosomes. To enrich post-synaptic densities (PSDs), the synaptosomal pellet was resuspended in Buffer B (0.32 M sucrose, 4 mM HEPES-NaOH, pH 7.4, 1% Triton X-100) and incubated on ice for 15 min, allowing selective solubilization of presynaptic membranes. The sample was then centrifuged at 13,000 × g for 20 min to separate the Triton-insoluble PSD fraction. The PSD pellet was washed and resuspended in Buffer C (50 mM HEPES-NaOH, pH 7.4, 2 mM EDTA, 0.5% Triton X-100) and subjected to a final centrifugation at 13,000 × g for 20 min (4 °C) to purify PSD components.

### Isolation of mitoEV-enriched fractions from primary astrocyte cultures

Conditioned cell culture medium (CM) was collected from astrocyte cultures 48h before EV isolation. CM was centrifuged at 500 *xg* for 15 min to remove living cells and at 2000 *xg* for 15 min to remove dead cells and larger apoptotic bodies. The supernatant was transferred and then run through the KrosFlo® Research IIi tangential flow filtration (TFF) system (Repligen, Spectrum Labs) with an mPES 300kD hollow fiber membrane (Repligen). The vesicles are simultaneously concentrated and diafiltered using 0.22 µm-filtered cold DPBS (Thermo Fisher Scientific). The final volume of 25 mL was transferred to a 38.5 mL, Open-Top Thinwall Ultra-Clear Tube (Beckman Coulter) and centrifuged at 100,000 *xg* for 70 min to pellet the EVs. The EV pellet was resuspended in 35 mL of DPBS and centrifuged at 100,000 *xg* for 70 min at 4°C to wash the EVs. To separate the different EV fractions, we used a high-resolution iodixanol step-gradient ultracentrifugation (modified from D’Acunzo *et al*.[15]). Briefly, the EV pellet was resuspended in 1.5 mL of an iodixanol solution (OptiPrep™, StemCell Technologies), layered in a 14 mL Open-Top Thinwall Ultra-Clear Tube (Beckman Coulter) and centrifuged at 200,000 *xg* for 16 h at 4°C. Afterwards, 8 fractions of 1.5 mL are collected from the top of the gradient. We pooled the small EV/exosome fractions (fractions 4–6) and did the same for the mitoEV-enriched fractions (fractions 7 and 8). Both were resuspended in DPBS and centrifuged at 100,000 *xg* for 70 min to wash out residual iodixanol. MitoEV pellets were stored at either 4°C for immediate use or -80°C for long-term storage.

### EV treatment in primary neurons

Crude EVs and mitoEV-enriched fractions were isolated from astrocyte-CM as described above, and protein concentration was quantified using a BCA assay. To deplete functional mitoEVs from the crude EV fractions, EVs were incubated with 2 µM carbonyl cyanide-4-(trifluoromethoxy)phenylhydrazone (FCCP; Sigma-Aldrich) for 30 min at 37 °C. Samples were washed once with PBS, then ultracentrifuged at 100,000 × g for 30 min. For crude EV treatments, neurons were exposed to EVs at a concentration corresponding to 0.5 ng EV protein per neuron for 6 h. For mitoEV treatments, a dose–response analysis was performed using increasing mitoEV concentrations (5.6, 12, 24, 48, and 96 pg protein per neuron), corresponding to a two-fold serial dilution, for 24 h.

### Nanoparticle trace analysis (NTA)

NTA was used to characterize the number and size of mitoEVs, and the analysis was performed with the NanoSight LM10-HSGF system (Malvern Panalytical). The NTA instrument is composed of a conventional light microscope combined with the NanoSight LM14C particle viewing unit, a 488 nm laser, a SCMOS camera, and a NanoSight Syringe Pump (Malvern Panalytical). The mitoEV samples were diluted in DPBS filtered through a 0.22 µm filter. The analysis was performed using NanoSight Software NTA 3.0 0060 (camera level 12–13, syringe pump speed 50, detection level 4), acquiring five videos of 20 s each, with 499 frames per sample.

### Cryogenic electron microscopy (cryo-EM)

Cryo-EM was used to analyze the structure of mitoEV fractions. R-23Cu mesh Quantifoil grids were glow-discharged (40mAm, 60 s) in a GloQube (Quorum). After isolation, the vesicles were resuspended in 30 μL EV-depleted Tris-HCl buffer (pH 7.2). Three μL of the EV solution was applied to the grid, blotted with filter paper (3-5 s) and then plunge-frozen in liquid ethane using a Vitrobot™ Mark IV (Thermo Fisher Scientific). Images were obtained with Talos Artica equipped with a Falcon 4i direct electron detector (DED) (Thermo Fisher Scientific) at 49000x magnification. Data were acquired with a total accumulated electron dose of ∼28 e⁻/Å² and a nominal defocus range of −3 to −6 µm.

### Transmission electron microscopy (TEM)

Three-month-old animals were anaesthetized and perfused with 2% glutaraldehyde and 1% formaldehyde in 0.1 M phosphate buffer solution via intracardiac perfusion. Brains were stored in fixing solution until hemispheres were cut in a brain slicer matrix and coronal slices collected for sectioning. Leica Ultracut UCT was used to create ultrathin sections, and uranyl acetate and lead citrate were used as contrasting agents. Sections were examined at 100 kV using a Tecnai 12 BioTWIN transmission electron microscope. Digital images were acquired from the hippocampus *Cornu Ammonis* area 1 (CA1) at a primary magnification of 30,000 x. Ten cells were snapped per brain, and 30-40 synapses were analyzed per condition. A synapse was considered when the presence of both synaptic vesicles and post-synaptic density was detected.

### Scanning electron microscopy (SEM)

Astrocyte–astrocyte co-cultures and astrocyte–neuron co-cultures were seeded onto glass coverslips. Cells were allowed to grow or differentiate for 5 days. Culture media were removed, and cells were rinsed twice with D-PBS, then fixed in 2.5% glutaraldehyde prepared in 0.1 M phosphate buffer for 1 h at room temperature. Samples were dehydrated through a graded ethanol series, dried, mounted on field-emission scanning electron microscopy (FESEM) stubs, and sputter-coated prior to imaging. High-resolution images of cell surface morphology were acquired using a Zeiss Ultra 55 field-emission scanning electron microscope. TNT length was quantified from SEM images using Fiji/ImageJ software.

### Mass spectrometry (MS) sample preparation and analysis

To eliminate iodixanol residue, isolated mitoEVs were briefly run on a 12% Bis-Tris gel (Invitrogen) in NuPAGE MES SDS running buffer (Thermo Fisher) until all proteins focused into a single 1-cm band. Gels were washed three times in ddH_2_O for 15 min each and visualized by staining with EZBlue™ Gel Staining Reagent (Sigma). Gel bands were excised, destained, dehydrated and rehydrated according to the protocol available in the Supplementary Methods. In-gel digestion was performed using a trypsin solution (25 ng/μL trypsin (MS grade, Sigma) in 10 mM AMBIC and 10% Acetonitrile), followed by 100 mM AMBIC overnight at 37°C. Using an extraction buffer (5% Formic Acid in 50% Acetonitrile), the supernatants containing the peptides were transferred to Lo-Bind Eppendorf tubes. Extractions were repeated and the pooled supernatants were concentrated in an Eppendorf Speedvac. The concentrated peptides were reconstituted in 30 μL of solution A* (2% acetonitrile, 1% trifluoroacetic acid) for analysis. The supernatant was carefully transferred and analyzed using a Nanodrop to determine peptide concentration.

Peptides were separated by nanoLC using the Thermo EasyLC 1200 HPLC system and analyzed on the Orbitrap Exploris (Thermo Fisher Scientific) run in DIA mode with FAIMS ProTM Interface (Thermo Fisher Scientific). Full MS spectra were collected across a m/z range of 400–1000. Detailed LC gradients and instrument parameters are provided in the Supplementary Methods. The Technical University of Denmark, through Biogenity, has contributed by performing the proteomics sample analysis.

### LC-MS Proteomic Data Analysis

RAW DIA-MS data were processed using SpectronautTM (version 19.1) against the *Mus musculus* UniProt database (taxonomy ID 10090). Downstream statistical and enrichment analyses were performed in R (v4.4.2) using Bioconductor 3.20 packages [26] [27]. Differential protein abundance was assessed using linear modeling with empirical Bayes moderation and FDR correction. Functional enrichment analyses (ORA and GSEA) were conducted using KEGG [28], Gene Ontology [29], Reactome [30], and custom mitochondrial annotation databases. Missing values were imputed and data were normalized prior to analysis (see Supplementary files for details). All code and processed data are available at https://github.com/jesusalarcongil/AstrocyteNeuronMitoEVs.

### Protein extracts and western blotting

Cells were lysed using RIPA buffer supplemented with protease and phosphatase inhibitors to obtain total cellular extracts. Subsequently, a benzonase solution (1 M Tris pH = 8.0, 0.5 M MgSO_4_ and 1x benzonase (Millipore) was used to degrade nucleic acids. The cell lysate was incubated on ice for 30 min, followed by centrifugation for 5 min at 14,000 *xg* at 4°C and the pellet, containing cell debris, was discarded. EVs and mitoEVs were lysed using supplemented RIPA buffer. Protein concentration was determined using the Pierce BCA Protein Assay Kit (Thermo Fisher). Samples were denatured in sample buffer for 5 min at 95 °C. 10-20 µg of protein were loaded into and separated on NuPAGE 4-12 % Bis-Tris gels (Invitrogen) in NuPAGE MES SDS running buffer. The ECL™ Rainbow™ Marker – Full Range (GE HealthCare) was used as the molecular marker. The proteins were transferred for 2 h to 0.45 µm nitrocellulose membranes (GE Healthcare), and membranes were incubated with Ponceau staining (Sigma-Aldrich) for 5 min to evaluate protein transfer. Afterwards, membranes were blocked for 1 h at RT with 5% BSA/TBS-T, then incubated overnight at 4°C with the primary antibodies listed in Supplementary Table S2, diluted in 5% BSA/TBS-T. β3-tubulin or Ponceau staining was used as the loading control. The membranes were washed and then incubated with the appropriate secondary antibodies listed in Supplementary Table S2, diluted in 1 % BSA/TBS-T, for 1 h at RT. The Odyssey CLx Imaging System (LI-COR Biosciences) was used to develop the membranes, and the image quantification was performed in Fiji. Synaptic cytosol fractions are normalized to synaptophysin; mitoEVs-enriched fractions are normalized to ponceau staining. Uncropped western blot images are provided in the Supplementary files.

### Immunocytochemistry and image analysis

Cells were fixed with 4% PFA in PBS for 15 min at RT at DIV10-12. When applied, before fixation, neurons are treated with CellTracker Blue. 0.2% Triton X-100 (Sigma-Aldrich) in PBS was added for 2 min to permeabilize the cells, followed by a 1 h blocking period with 3 % Bovine Serum Albumin (BSA)/PBS (Sigma-Aldrich). The cells were incubated overnight at 4°C with the primary antibody TOM20 (1:200, Santa Cruz Biotechnology) diluted in 3% BSA/PBS. After three washing steps, the cells were incubated for 1 h at RT with the appropriate Alexa Fluor secondary antibodies (1:500, 594 anti-rabbit, Invitrogen) diluted in 3 % BSA/PBS. To stain TNTs, phalloidin-633 probe (Thermo Fisher) was used in combination with secondary antibodies. Nuclei were stained with a 4 µg/mL Hoechst solution prepared in PBS and incubated for 15 min at RT. Lastly, the Antifade mounting media (VectaShield Plus) was used to mount the coverslips. Cells are imaged with a Nikon CrEST X Light V3 Spinning Disk microscope using a 60X objective. For mitochondrial transfer quantification, z-stack volumes with a 0.2 µm step size were acquired. Astrocyte-to-neuron mitochondrial transfer was quantified using Imaris. Three surfaces are created for the CellTracker Blue, the mEmerald-mito and the mitoDsred signals. MitoDsred surfaces with a bigger volume than a sphere with a 1 µm radius are excluded, to keep only mitoEV-like structures. The overlapping ratio between mitoDsred surface with CellTracker Blue and mEmerald-mito surfaces is used to quantify integration in the host cell and the host cell mitochondrial network, respectively. In astrocytes, the total number of mitoDsRed mitoEVs was counted and classified into three different subgroups – integrated (full overlap with TOM20), attached (partial overlap with TOM20), and free. For the co-localization analysis of mitochondria with synaptophysin, Mander’s M2 coefficient was applied.

### Mitophagy events assessment in astrocytes

After 48 h of co-culture with neurons, astrocytes transfected with mt-Keima were washed and incubated in 1 x ACSF supplemented with 1.5 mM CaCl_2_ and 1.5 mM MgCl_2_. Live imaging was performed at 37 °C to assess the number of mitophagy events in astrocytes. The z-stack images were taken on the LSM980-Airy confocal microscope using a 63x lens with NA=1.4%. Fluorescence was detected by using two different excitation wavelengths – 458 nm (green) and 561 nm (red) – and emission at 620 nm. The Fiji software was used to analyze the obtained images. Z-stack images were processed, and ROIs were designed in the resultant 3D reconstructions. A threshold was applied, and fluorescence was measured in both green and red channels. The ratio of the % area of red to green signal indicates the % of mitochondria localizing within the lysosome.

### Seahorse extracellular flux analysis

Mitochondrial respiration and glycolytic function were assessed using a Seahorse XF96 extracellular flux analyzer (Agilent) to measure oxygen consumption rate (OCR) and extracellular acidification rate (ECAR). Astrocytes, neurons, and isolated mitoEVs were analyzed. Standard mitochondrial and glycolytic stress tests were performed using sequential injections of oligomycin, FCCP, and rotenone/antimycin A. Fatty acid oxidation (FAO)-dependent respiration in neurons was assessed using palmitate–BSA substrate conditions. MitoEV respiratory capacity was evaluated under substrate-driven conditions to assess intrinsic electron transport chain activity, as previously described [31]. OCR and ECAR values were normalized to protein content. Detailed experimental conditions and reagent concentrations are provided in the Supplementary files.

### Lipid droplet staining

Cells were simultaneously stained with 5 μM BODIPY™ 493/503 (Thermo Fisher) to label neutral lipids in lipid droplets and 10 μM CellTracker™ Red CMTPX (Thermo Fisher) to identify individual neurons. Staining was performed directly in half-volume conditioned neuronal medium, followed by a 20-min incubation at 37 °C. After incubation, the dye-containing medium was removed, and cells were washed twice before imaging in Phenol Red–free Neurobasal medium (Thermo Fisher). Imaging was performed on a Zeiss LSM980 with Airyscan using a 40× water-immersion objective. Z-stack images were acquired and processed in Fiji to generate 3D reconstructions. The lipid droplet area within each ROI was quantified using the Analyze Particles function in Fiji, and mean values were calculated. Data were normalized to the number of neurons.

### Statistical analysis

Statistical analysis was performed using GraphPad Prism (version 9). Data were presented as mean ± SEM and n refers to the number of unique individual biological samples. No blinding nor randomization of samples was performed. Data distribution was assessed using normality tests and log-transformations were applied when appropriate. To make comparisons between groups (genotype or treatments), non-parametric Kruskal-Wallis test was used, followed by Dunn’s multiple comparison test, or unpaired t-test (two-tailed). For normal distributions, one-way ANOVA followed by post-hoc Šídák’s multiple comparison test was performed. Data obtained from live imaging analysis in astrocytes were analyzed with an unpaired t-test for normal and homoscedastic data, a Welch’s test for normal and heteroscedastic data, or a Kolmogorov-Smirnov test for non-normal data, with a p < 0.05 being considered significant. Outliers were detected using the ROUT method (Q=1%).

## RESULTS

### *App^NL-G-F^* neurons exhibit presynaptic mitochondrial loss and impaired synaptic energetics

Presynaptic terminals require a constant local supply of ATP and depend on efficient mitochondrial positioning at synapses [32]. We have previously reported that late symptomatic *App^NL-G-F^* mice (10-12 months old), a well-characterized model that recapitulates AD-associated Aβ pathology, display abnormal synaptic organization associated with deficient autophagy and mitochondrial loss [4]. In contrast, at early stages, the *App^NL-G-F^* model exhibits a compensatory increase in mitochondrial activity [4]. Here, we asked whether, despite increased mitochondrial function, early symptomatic *App^NL-G-F^* mice exhibit a localized loss of presynaptic mitochondria that could influence synaptic function. We observed a significant depletion of mitochondria in the presynaptic terminals of the CA1 region of the hippocampus in *App^NL-G-F^* mice as early as 3 months of age (p = 0.0111) (**Fig 1a**), preceding the appearance of Aβ deposits in this region [4, 23]. To further examine presynaptic mitochondria distribution, we analyzed synaptosome fractions isolated from primary neuronal cultures derived from *App^NL-G-F^* mice (**Fig 1b, c**). No changes were detected in the levels of key mitochondrial proteins, including TOM20 (import), MCU (calcium uptake), and PDH (linking glycolysis to the tricarboxylic acid cycle, TCA) (**Fig 1c, Supplementary Fig S1a**). In contrast, colocalization analysis of the F_1_F_0_-ATP synthase subunit *c* (Complex V) with the synaptic vesicle protein synaptophysin revealed a reduced mitochondrial presence in *App^NL-G-F^* axonal processes (p = 0.0066) (**Fig 1d, Supplementary Fig S1b**).

**Figure 1:**
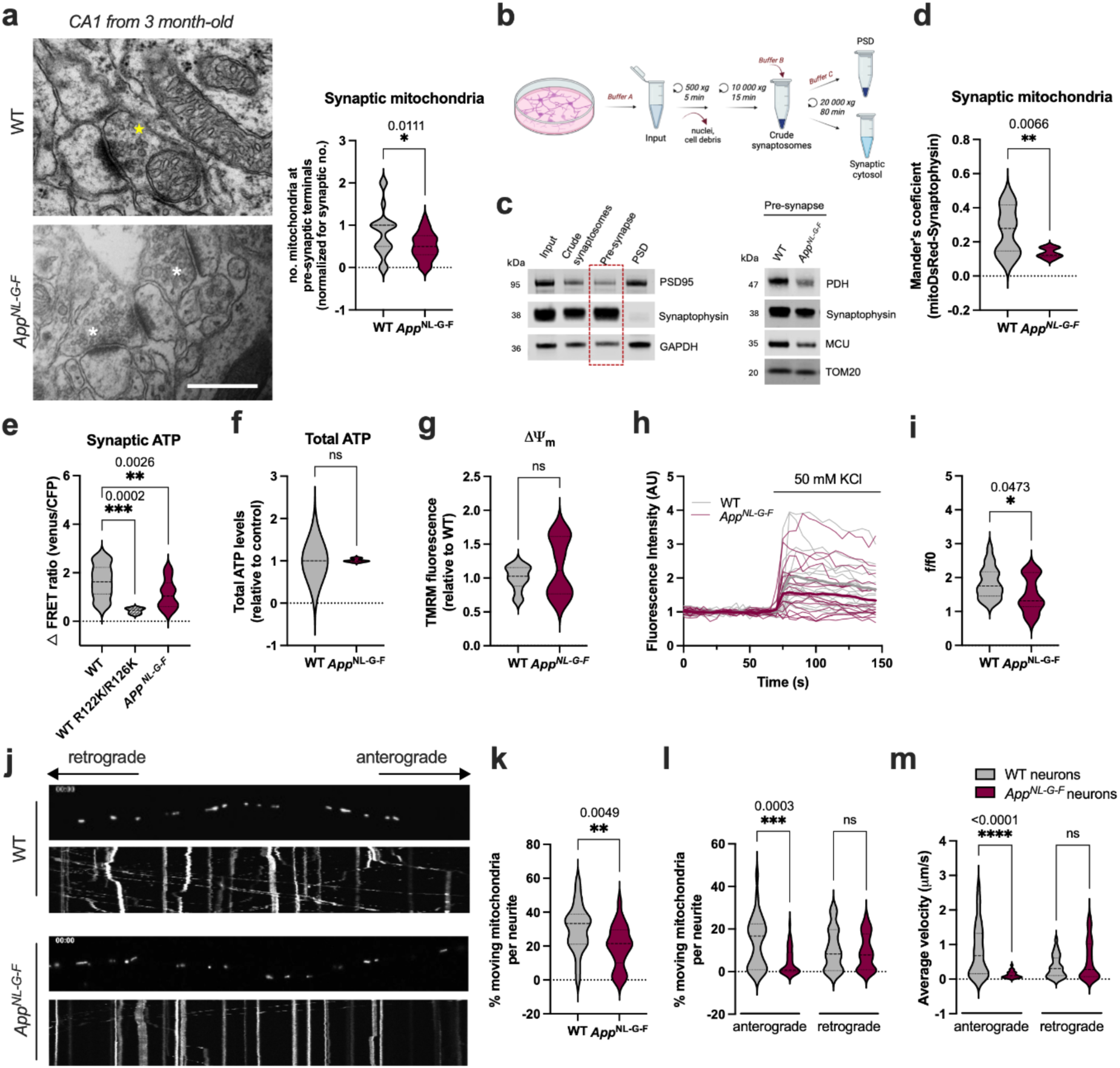
**Mitochondrial anterograde transport and synaptic ATP levels are affected in App knock-in neurons**. **(a)** Representative TEM images showing mitochondria at the presynaptic terminal in WT and *App^NL-G-F^*mice hippocampus. Asterisk indicates presynaptic terminals containing synaptic vesicles, yellow: with mitochondria, white: without mitochondria. Scale bar = 500 nm. Quantification of the number of mitochondria at the pre-synapse was normalized to the number of synapses (n = 27-40 synapses from 4 mice per genotype). **(b)** Schematic representation of synaptosome isolation and enrichment of the pre-synapse (synaptic cytosol) and post-synaptic density (PSD) from primary neuron cultures. **(c)** Representative Western blots showing the purity of the fractions and mitochondrial proteins in WT and *App^NL-G-F^* synaptic cytosolic fractions. Quantification is shown in Fig S1a. **(d)** Colocalization analysis of synaptophysin and subunit *c* of mitochondrial F_1_F_0_-ATP synthase (in mitoDsRed-transfected neurons) (n = 12 from 3 independent cultures). **(e)** Quantification of synaptic ATP using the FRET-based ATP sensor SypI-ATeam1.03. WT R122K/R126K carry a F_0_F_1_ ATPase subunit insensitive to ATP. **(f)** Quantification of total ATP levels in neuron homogenates (n = 3). **(g)** Quantification of mitochondrial membrane potential (Δψ_m_) in neurons using TMRM fluorescent probe (10 nM) (n = 15 from 3 independent cultures). **(h)** Traces of fluorescent levels of SynaptopHluorin, directly proportional to the synaptic vesicle release responsive to KCl. **(i)** Quantification of the total synaptic vesicle pool released by KCl stimulation for WT and *App^NL-G-F^*neurons (n = 16-19, from 3 independent cultures). **(j)** Representative kymographs (xx, distance *vs* yy, time) of mitochondrial transport in mitoDsRed-transfected neurons. **(k-m)** Percentage of moving mitochondria, direction, and velocity of movement were quantified through kymographs (n = 26-27 neurites from 4 independent cultures). Data represented as mean ± SEM. Statistical significance was analyzed using an unpaired t-test. *p < 0.05, **p < 0.01, ***p < 0.001, ****p < 0.0001.

To assess synaptic ATP levels, we used a synaptic targeted Förster Resonance Energy Transfer (FRET)-based ATP sensor. This construct reports ATP binding through changes in FRET efficiency and is targeted to presynaptic terminals via synaptophysin [24]. Sensor specificity was validated using an ATP-insensitive mutant (**Fig 1e**). Using this approach, we found a significant reduction in synaptic ATP levels in *App^NL-G-F^* neurons (p = 0.0026), whereas total ATP levels remained unchanged (**Fig 1e, f**). Consistently, mitochondrial membrane potential (ι14′_m_) was not altered (**Fig 1g, Supplementary Fig S1c**), suggesting the reduction in synaptic ATP at these early stages is unlikely to result from gross defects in the mitochondrial electron transport chain but rather a local impairment at synaptic sites. Synaptic vesicle release is regulated by Ca^2+^ and ATP [33], and thus reduced synaptic ATP availability may compromise synaptic function. We therefore used the synaptophysin-pHluorin (SyPhy) reporter to examine changes in synaptic vesicle release in *App^NL-G-F^* neurons. Following 1 min of baseline recording, neurons were depolarized by the addition of 50 mM KCl to induce synaptic vesicle exocytosis. We observed a mild, but significant reduction in synaptic vesicle release in *App^NL-G-F^* primary neurons compared to WT controls (p = 0.0473) (**Fig 1h, I, Supplementary Fig S1d**).

We next asked whether compromised mitochondrial anterograde transport could underlie the reduction in presynaptic mitochondrial pool observed in early-symptomatic *App^NL-G-F^* neurons. Mitochondrial movement and velocity along the axons were examined using kymograph analysis. This revealed a significant reduction in anterograde, but not retrograde, axonal transport in *App^NL-G-F^* neurons (p = 0.003) (**Fig 1j-m**). In addition, mitochondrial mean velocity was markedly decreased (p < 0.0001) (**Fig 1m**), indicating that defective axonal transport is a major driver of presynaptic mitochondrial depletion.

Taken together, these data indicate that impaired axonal transport and a reduced number of presynaptic mitochondria may cause a significant loss of synaptic ATP, alter synaptic vesicle release and ultimately lead to synaptic impairment in *App^NL-G-F^*neurons.

### AD astrocytes exhibit a less complex but more dynamic mitochondrial network

Astrocytes play a central role in brain homeostasis and metabolic support to the neurons [34]; yet the dynamics of their mitochondrial network – particularly in live cells and in AD – remain poorly defined. To investigate this, WT and *App^NL-G-F^* astrocytes were transduced with mitoDsRed and analyzed by 4D spinning-disk confocal life-cell imaging to assess mitochondrial network connectivity and dynamics (**Fig 2a, b**). Deconvolved images were computationally segmented using MitoGraph [35] and analyzed with mitoTNT (Mitochondrial Temporal Network Tracking) algorithm to extract morphological and temporal network parameters (**Fig 2a, b**).

**Figure 2:**
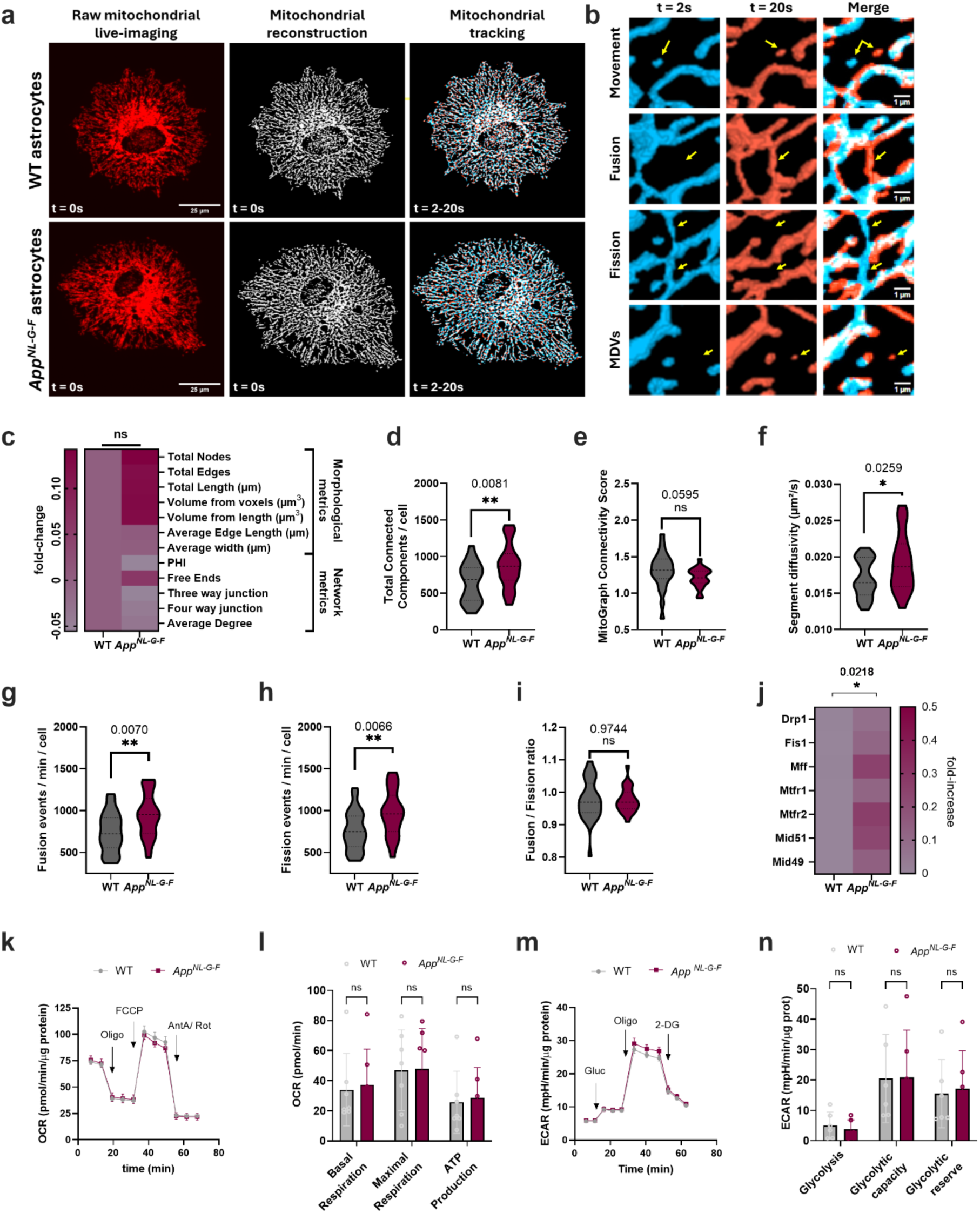
Altered mitochondrial dynamics in primary astrocytes. **(a)** Pipeline of imaging analysis of 4D mitochondrial network time-lapses in mitoDsRed-infected astrocytes. Maximum projection images of mitochondrial segmentation (second column) show the surface of the mitochondrial network in white. The third column shows the tracking between two time points (2 s, in blue; 20 s in red) and static mitochondrial voxels in white. **(b)** Representative images of mitochondrial movement, fusion, fission, and MDVs detected between the 2 s time point (blue) and the 20 s time point (red). **(c-i)** Mitochondrial parameters including **(c)** heatmap of mitochondrial morphology and network metrics, **(d)** total number of mitochondrial connected components by cell, **(e)** MitoGraph Connectivity Score, **(f)** mitochondrial segment diffusivity, **(g-i)** fission and fusion events (n = 25 images from 3 independent cultures). **(j)** Heatmap presents the expression of fission-related genes in WT and *App^NL-G-F^*primary astrocytes (n = 5). **(k, l)** Seahorse analysis of OCR in WT and *App^NL-G-F^* primary astrocytes following sequential injection of oligomycin (Oligo), FCCP, and rotenone + antimycin A (Rot + Ant A), as shown by representative traces of OCR. Bar graph shows quantification of basal and maximal respiration, and ATP production. Data shown as mean ± SEM (n = 7). **(m, n)** Seahorse analysis of extracellular acidification rate (ECAR) in WT and *App^NL-G-F^*primary astrocytes following sequential injection of glucose (Gluc), oligomycin (Oligo) and 2-deoxy-D-glucose (2-DG), as shown by representative traces of ECAR. Bar graph shows quantification of glycolysis, glycolytic capacity and reserve. Data shown as mean ± SEM (n = 6). Statistical significance was analyzed using an unpaired t-test, Mann-Whitney test (g, h) or 2-way ANOVA (in k, m). * p ≤ 0.05, ns. non-significant.

Quantification of mitochondrial morphology revealed no significant differences in volume or length between WT and *App^NL-G-F^*astrocytes, despite a mild upward trend in *App^NL-G-F^* (**Fig 2c**). Similarly, most mitochondrial network metrics remain unchanged, with only a trend toward lower mitochondrial complexity in *App^NL-G-F^* astrocytes (**Fig 2c**). Nevertheless, the *App^NL-G-F^*astrocytic network exhibits a significant increase in independent mitochondrial components (p = 0.0081) and a reduced connectivity as measured by the MitoGraph Connectivity Score (p = 0.0595) (**Fig 2d, e**). To assess mitochondrial network motility, we quantified mitochondrial segment diffusivity, which was significantly increased in *App^NL-G-F^* astrocytes compared with WT astrocytes (p = 0.0259), indicating enhanced movement (**Fig 2f**). In parallel, analysis of mitochondrial fusion and fission events revealed a significant increase in both fusion (p = 0.0070) and fission events (p = 0.0066) in *App^NL-G-F^* astrocytes, resulting in an overall unchanged fusion-fission balance (**Fig 2g-i**). In agreement, an overall increase in the expression levels of fission proteins was observed in *App^NL-G-F^* astrocytes compared with WT controls (p = 0.0218 for genotype) (**Fig 2j**), which corroborates reduced connectivity. Together, these findings indicate that the mitochondrial network in *App^NL-G-F^* astrocytes is less interconnected and more dynamic, showing more remodeling events.

To determine whether the observed alterations in mitochondrial network dynamics were associated with changes in astrocytic bioenergetics, we assessed mitochondrial respiration and glycolytic function using Seahorse extracellular flux analysis. No significant differences were observed between WT and *App^NL-G-F^* astrocytes in basal or maximal mitochondrial respiration, complex V–linked ATP production, glycolytic activity, or glycolytic reserve **(Fig 2k–n)**. Together, these findings show that mitochondrial network remodeling in *App^NL-G-F^*astrocytes occurs despite preserved astrocytic bioenergetic function at this stage of the disease.

### Astrocyte-to-neuron mitochondrial transfer occurs at axons and is partially mediated by mitoEVs

Given the early loss of presynaptic mitochondria in *App^NL-G-F^*neurons (**Fig 1**), and the increased mitochondrial remodeling and dynamics observed in astrocytes at the same disease stage (**Fig 2**), we hypothesized that astrocytes may engage compensatory mechanisms to support neuronal bioenergetic demand through intercellular mitochondrial transfer. Earlier findings show that astrocytes can improve neuronal function by transferring healthy mitochondria to stressed neighboring neurons [11]. This prompted us to investigate whether mitochondrial exchange between astrocytes and neurons can occur locally at axonal compartments and whether this process is altered in the *App^NL-G-F^*model.

Using microfluidic chambers that physically separate neuronal somata from distal axons, we assessed whether astrocyte-derived mitochondria can be transferred locally to axonal compartments (**Fig 3a**). Neuronal arborization was labelled with EGFP under the Synapsin I promoter and astrocytic mitochondrial network labelled by transducing cells with mitoDsRed under the GFAP promoter. In this system, WT neuronal axons extended through 400 μm-long microgrooves to contact astrocytes cultured in the distal compartment, allowing axon-specific interactions in the absence of soma-derived signals. High-resolution imaging revealed the presence of astrocyte-derived mitochondria within the axonal network, which appeared predominantly as small, punctate structures (**Fig 3b**). These data demonstrate that local axonal communication with astrocytes is sufficient to support mitochondrial transfer.

**Figure 3:**
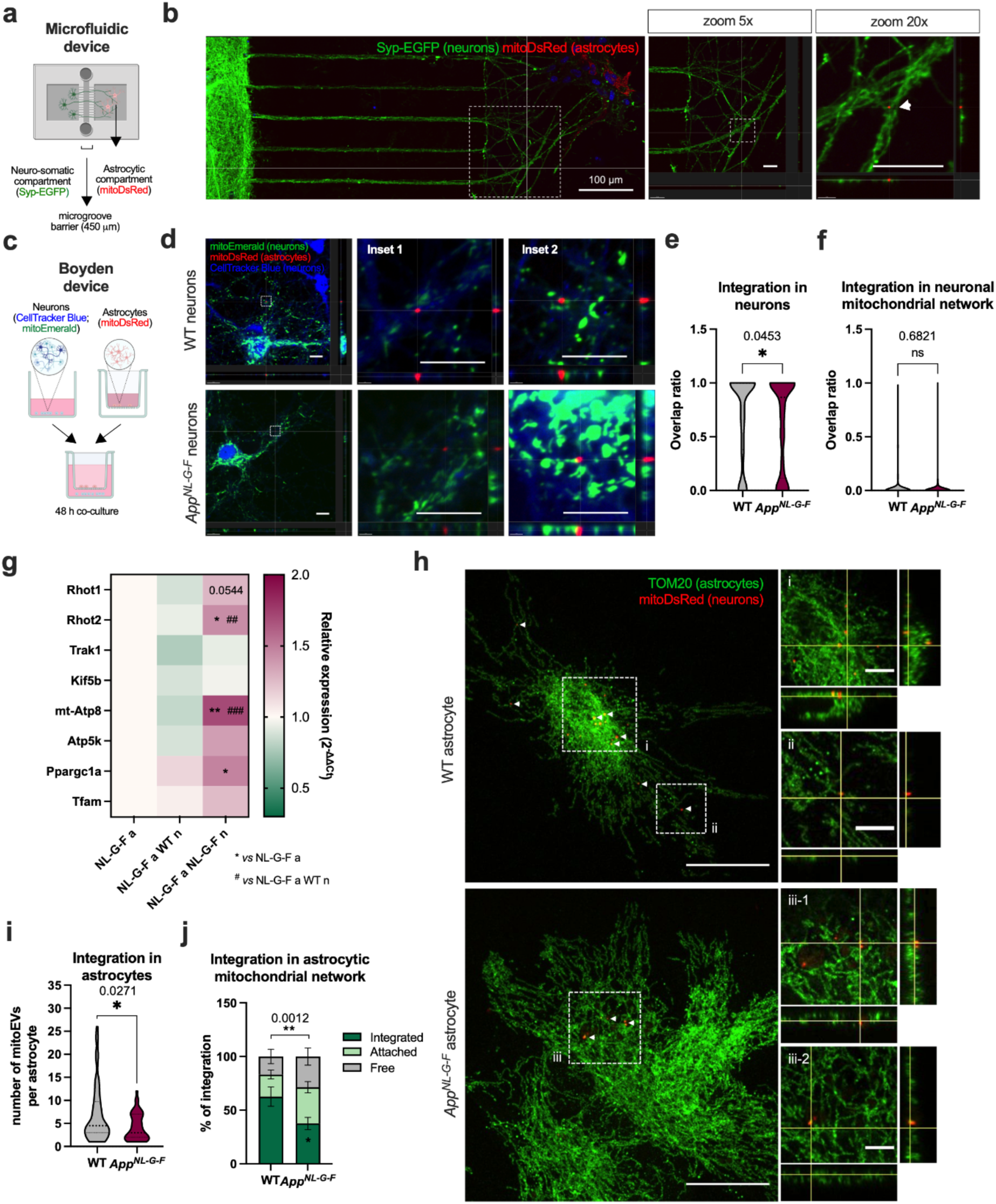
Bidirectional mitochondrial transfer in neuron-astrocyte co-cultures. Co-cultures were established in **(a, b)** microfluidic chambers to allow direct cell communication between axons and astrocytes. Neurons plated in microfluid chambers with 400 μm-long microgrooves were transduced with Syp-GFP, and astrocytes with mitoDsRed to differentially label axons in green and astrocytic mitochondria in red. Scale bar = 100 µm; insets = 20 µm. **(c-f)** Boyden chambers were used to allow defined cell-free gaps between neurons and astrocytes. Host neurons transduced with mitoEmerald were plated in the outer chamber, donor astrocytes transduced with mitoDsRed were plated in the inner chamber. Neurons were labeled with CellTracker Blue. Scale bar = 10 µm. **(e)** Mitochondrial integration in neurons and in the **(f)** neuronal mitochondrial network were analyzed and quantified using Imaris software. **(g)** Heatmap shows the expression levels of mitochondrial transport, OXPHOS, and mitochondrial biogenesis-related genes in *App^NL-G-F^* astrocytes, in the absence or presence of neurons in co-culture, in Boyden chambers. * p ≤ 0.05, ** p ≤ 0.01 *versus* astrocytes in monoculture; ^##^ p ≤ 0.01, ^###^ p ≤ 0.001 *versus* astrocytes in co-culture with WT neurons. **(h)** Representative confocal images of neuron-to-astrocyte mitochondrial transfer; neurons were transduced with MitoDsRed and astrocytes labelled with TOM20. Scale bars = 20 µm/ inset = 5 µm. Quantification of **(i)** mitochondrial integration in astrocytes and in the **(j)** astrocytic mitochondrial network. Data presented as mean ± SEM. * p ≤ 0.05, ** p ≤ 0.01.

Given the predominantly small punctate morphology of astrocyte-derived mitochondrial structures observed at axonal sites, we next sought to determine whether astrocyte-to-neuron mitochondrial transfer occurs via vesicle-mediated mechanisms. To this end, we set up co-cultures in Boyden chambers to restrict intercellular communication to diffusible factors and EVs (**Fig 3c**). Exposure of astrocytes to neuronal culture medium did not significantly affect astrocyte or mitochondrial morphology (**Supplementary Fig S2a-c**), excluding media-induced stress effects. High-resolution confocal imaging revealed astrocyte-derived mitochondrial puncta overlapping with CellTracker Blue-labeled neurons, consistent with uptake of mitoEVs (**Fig 3d**). Quantitative analysis showed that astrocyte-to-neuron mitochondrial transfer was significantly reduced in *App^NL-G-F^* co-cultures compared with WT (p = 0.0453) (**Fig 3e**). To assess whether transferred astrocytic mitoEVs integrate into the neuronal mitochondrial network, we performed colocalization analysis between astrocyte-derived and neuronal mitochondrial signals. This analysis revealed that mitoEV integration into the neuronal mitochondrial network was limited and did not differ between genotypes (p = 0.6821) (**Fig 3f**). Because tunneling nanotubes (TNTs) have also been implicated in mitochondrial transfer[36], we next assessed TNT formation using scanning electron microscopy. TNTs were readily observed between astrocytes (**Supplementary Fig S2d**), with *App^NL-G-F^* astrocytes exhibiting a higher proportion of shorter TNTs (40–60 µm) compared with WT astrocytes (**Supplementary Fig S2e**). In contrast, under our co-culture conditions astrocyte-to-neuron TNT formation was not detected. We cannot exclude technical limitations associated with the highly arborized neuronal morphology and the lack of specific TNT markers, which hinder the unambiguous distinction between neuronal processes and TNTs. Despite these structural differences, rotenone treatment induced comparable mitochondrial transfer between astrocytes in WT and *App^NL-G-F^*cultures (p = 0.1102) (**Supplementary Fig S2f, g**). Together, these findings indicate that, in our system, astrocyte-to-neuron mitochondrial exchange occurs predominantly via vesicle-mediated mechanisms rather than TNTs.

To determine whether *App^NL-G-F^* astrocytes transcriptionally adapt their mitochondrial machinery in response to neuronal cues, we next analyzed the expression of mitochondria-associated genes in astrocytes cultured alone or co-cultured with neurons (**Fig 3c, g**). Co-culture with WT neurons did not induce significant transcriptional changes, whereas astrocytes exposed to *App^NL-G-F^*neurons upregulated the expression of genes associated with mitochondrial transport, MDV/mitoEV tubulation (e.g., Rhot1, Rhot2) [17, 37] and mitochondrial biogenesis (e.g., mt-Atp8, Ppargc1a) (**Fig 3g**). This response likely reflects an attempt to modulate mitochondrial distribution and availability under pathological conditions.

Together, these data indicate that astrocyte-to-neuron mitochondrial transfer occurs at axonal sites via vesicle-mediated mechanism and is impaired in the *App^NL-G-F^* model despite transcriptional adaptation of astrocytic mitochondrial pathways.

### Neuron-to-astrocyte mitochondrial transfer and fate in astrocytes

To determine whether mitochondrial transfer is bidirectional, we established a reverse Boyden chamber configuration and assessed neuron-to-astrocyte mitochondrial transfer. In this setup, neuronal mitochondria were fluorescently labelled with mitoDsRed. At the same time, the astrocytic mitochondrial network was identified by labeling the mitochondrial outer membrane (MMOM) protein TOM20 (**Fig 3h**). High-resolution imaging revealed astrocyte-associated neuronal mitoEVs at different stages of integration within the astrocytic mitochondrial network, including fully integrated, attached, and non-associated free structures (**Fig 3h**). Quantitative analysis showed that the number of mitoEVs transferred from neurons to astrocytes was significantly reduced in *App^NL-G-F^* co-cultures compared with WT (p = 0.0271) (**Fig 3i**). Additionally, in WT conditions, more than 50% of neuron-derived mitoEVs became fully integrated into the astrocytic mitochondrial network. This proportion was significantly reduced in *App^NL-G-F^* co-cultures (p = 0.0012), with a corresponding increase in attached but non-integrated mitoEVs (**Fig 3j**). As these analyses were performed on fixed samples, the observed mitochondrial association states represent static snapshots that may precede complete integration of transferred mitoEVs.

To investigate the fate of non-integrated mitoEVs in astrocytes, we assessed mitophagy in astrocytes in co-culture with neurons. Astrocytes co-cultured with *App^NL-G-F^* neurons exhibited increased expression of genes involved in both Parkin/PINK1-dependent (e.g., *Pink1, Parkin*) and -independent (e.g., *Fundc1, Ambra1*) mitophagy pathways, which was not evident in WT co-cultures (**Supplementary Fig S3a**). In addition, this was accompanied by increased mitophagy events (p = 0.0367), evidenced by elevated mt-Keima red puncta (mitochondria engulfed in lysosomes) (**Supplementary Fig S3b**). These findings reveal an asymmetry in intercellular mitoEV exchange between neurons and astrocytes and support a model in which neuron-derived mitoEVs may be preferentially routed toward astrocytic mitophagy in AD, consistent with enhanced transmitophagy[38, 39]. This aligns with our previous observations of autophagic impairment in axonal terminals in this knock-in AD model[4], suggesting that increased transfer of neuronal mitochondria to astrocytes may represent a compensatory mechanism to maintain mitochondrial quality control when it is impaired in the neurons.

### Astrocytic mitoEVs exhibit a bioenergetic and metabolic proteomic signature

To better understand the molecular and functional aspects of astrocyte-derived mitoEVs, we performed a comprehensive structural, proteomic and functional characterization. We established a protocol to isolate the total EV pool from the conditioned media of primary astrocytes in culture using TFF to concentrate conditioned media and remove soluble contaminants, followed by ultracentrifugation with a high-resolution iodixanol density gradient to separate EV subpopulations (modified from D’Acunzo *et al.*[15, 16]) (**Fig 4a**). Sequential fractions enriched for small EVs (sEVs; F4-6), and mitoEVs (F7-8) were isolated and analyzed separately. Immunoblot analysis confirmed that F4-6 were enriched for canonical sEV markers Alix and Flotilin-1, while F7-8 were enriched for mitochondrial proteins including the MOM transporter VDAC, the mitochondrial matrix protein pyruvate dehydrogenase PdhE1α, and mitochondrial inner membrane (MIM) subunits of the mitochondrial respiratory Complexes II and III (Sdhb and Uqcrc2, respectively). Additionally, we confirmed the absence of TOM20 (**Fig 4b**). We next quantified the number and size of the mitoEVs secreted by WT and *App^NL-G-F^* astrocytes using NTA, which revealed comparable particle numbers and size distributions between genotypes (**Fig 4c-e**), contradicting previous data suggesting that stress and disease conditions enhance EVs secretion [22, 40]. These findings indicate that the decreased mitochondrial transfer observed in *App^NL-G-F^* cultures is unlikely to result from reduced mitoEV release. In contrast, cryo-EM analysis revealed a significant reduction in mitoEV size in *App^NL-G-F^* astrocytes (p < 0.0001) (**Fig 4f, g**), suggesting that changes in mitoEV ultrastructure may not be fully captured by bulk particle analysis.

**Figure 4.**
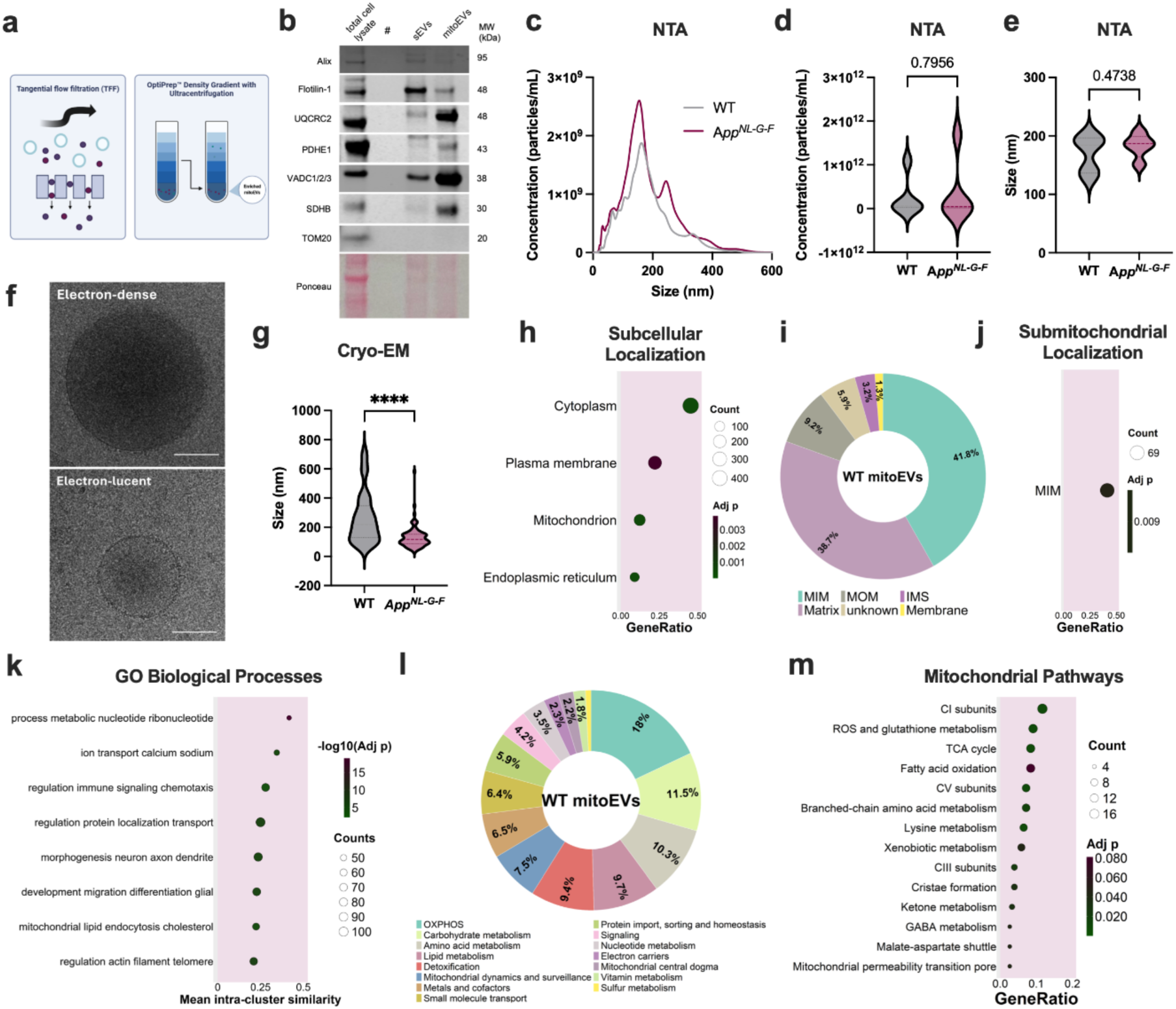
Structural, molecular and functional profile of astrocytic mitoEVs. **(a)** Experimental setup for mitoEVs isolation from conditioned medium. **(b)** Representative Western blot for sEV versus mitoEVs markers. **(c)** Vesicle size and concentration distribution (mitoEVs, WT vs *App^NL-G-F^*) measured with NTA. Quantification of **(d)** concentration and **(e)** size of mitoEVs by NTA. **(f)** Representative cryo-genic electron micrographs showing size and ultrastructure of WT mitoEVs (n=36-43 micrographs analyzed from 2 independent isolations). Scale bar = 100 nm. **(g)** Quantifications of vesicle size by cryo-EM. **(h)** ORA dotplot for subcellular location terms. **(i)** Doughnut chart showing the relative intensity percentages of proteins associated with submitochondrial location terms in WT mitoEVs. **(j)** ORA dotplot for submitochondrial location terms. **(k)** Dotplot of over-represented GO BP term clusters in WT mitoEVs. **(l)** Doughnut chart of mitochondrial pathway terms in WT mitoEVs. **(m)** ORA dotplot for mitochondrial pathways subterms for detected proteins in WT mitoEVs.

Contrary to sEVs, mitoEVs have been described to be highly enriched in protein content[16]. To further characterize the proteomic composition of astrocyte-derived mitoEVs, we performed label-free data-independent acquisition tandem mass spectrometry (DIA-MS/MS), allowing for an unbiased and comprehensive analysis of vesicle-associated proteins. A total of 1,305 proteins were identified, with 98.7% detected in both WT- and *App^NL-G-F^*-derived mitoEV samples (1,288 proteins) (**Supplementary Fig S4a**), indicating a largely conserved proteome between genotypes. Proteomic annotation using standard GO Cellular Component (GO CC) analysis combined with custom subcellular localization revealed that mitoEV-enriched fractions contained proteins annotated to multiple cellular compartments (**Supplementary Fig S4b, c**). Over-Representation Analysis (ORA) further demonstrated a significant enrichment of proteins originating from the cytoplasm, plasma membrane, mitochondria, and endoplasmic reticulum (ER) (**Fig 4h**). Within the mitochondrial subset, proteins were predominantly localized to the MIM (41.8%), matrix (38.7%), and MOM (9.2%) (**Fig 4i**), with MIM proteins being significantly overrepresented (**Fig 4j**).

To explore the functional landscape of the WT mitoEV proteome, we performed pathway enrichment analysis using KEGG, GO Biological Processes (GO BP), and custom mitochondrial pathway annotations. KEGG clusters revealed enrichment of metabolic pathways related to carbohydrate, amino acid, and lipid metabolism, as well as processes linked to the nervous and endocrine systems, and human diseases, including neurodegenerative diseases **(Supplementary Fig S4c)**. GO BP terms further highlighted processes such as nucleotide metabolism and the storage and transport of organic molecules, including lipids, amino acids, and carbohydrates **(Fig 4k)**. Mitochondrial pathway analysis revealed a predominance of proteins associated with OXPHOS (18%), carbohydrate metabolism (11.5%), amino acid metabolism (10.3%), lipid metabolism (9.7%), and detoxification (9.4%), along with pathways involved in mitochondrial dynamics and homeostasis (**Fig 4l, Supplementary Fig S4d**). Further analysis identified enrichments in functionally relevant subcategories, including selective respiratory complexes (CI, CIII, and CV), redox and glutathione metabolism, the TCA cycle, branched-chain amino acid and lysine metabolism, ketone metabolism, and FAO pathways (**Fig 4m, Supplementary Fig 4f, g**).

Taken together, these analyses establish that astrocyte-derived mitoEVs are enriched in proteins functionally linked to mitochondrial bioenergetics, intermediary metabolism, and cellular detoxification, providing a reference framework for subsequent comparison with the *App^NL-G-F^* mitoEV proteome.

### *App* knock-in-derived mitoEVs are depleted in mitochondrial proteins and exhibit impaired respiratory function

To identify disease-associated alterations in mitoEV composition, we compared *App^NL-G-F^* and WT mitoEV proteomes using a hybrid analytical approach (see Methods for details). Principal Component Analysis (PCA) revealed a clear separation between WT and *App^NL-G-F^* mitoEVs, with the first two components explaining more than 35% of the variance (**Fig 5a**), indicating distinct proteomic signatures. Gene Set Enrichment Analyses (GSEA) on custom annotations revealed a significant depletion of mitochondrial proteins in *App^NL-G-F^* versus WT mitoEVs **(Fig 5b**), driven primarily by the loss of proteins localized to the MIM and matrix (**Fig 5c**). Furthermore, OXPHOS- and lipid metabolism-related proteins showed a trend toward reduced abundance in WT and *App^NL-G-F^* mitoEVs (**Fig 5d**).

**Figure 5.**
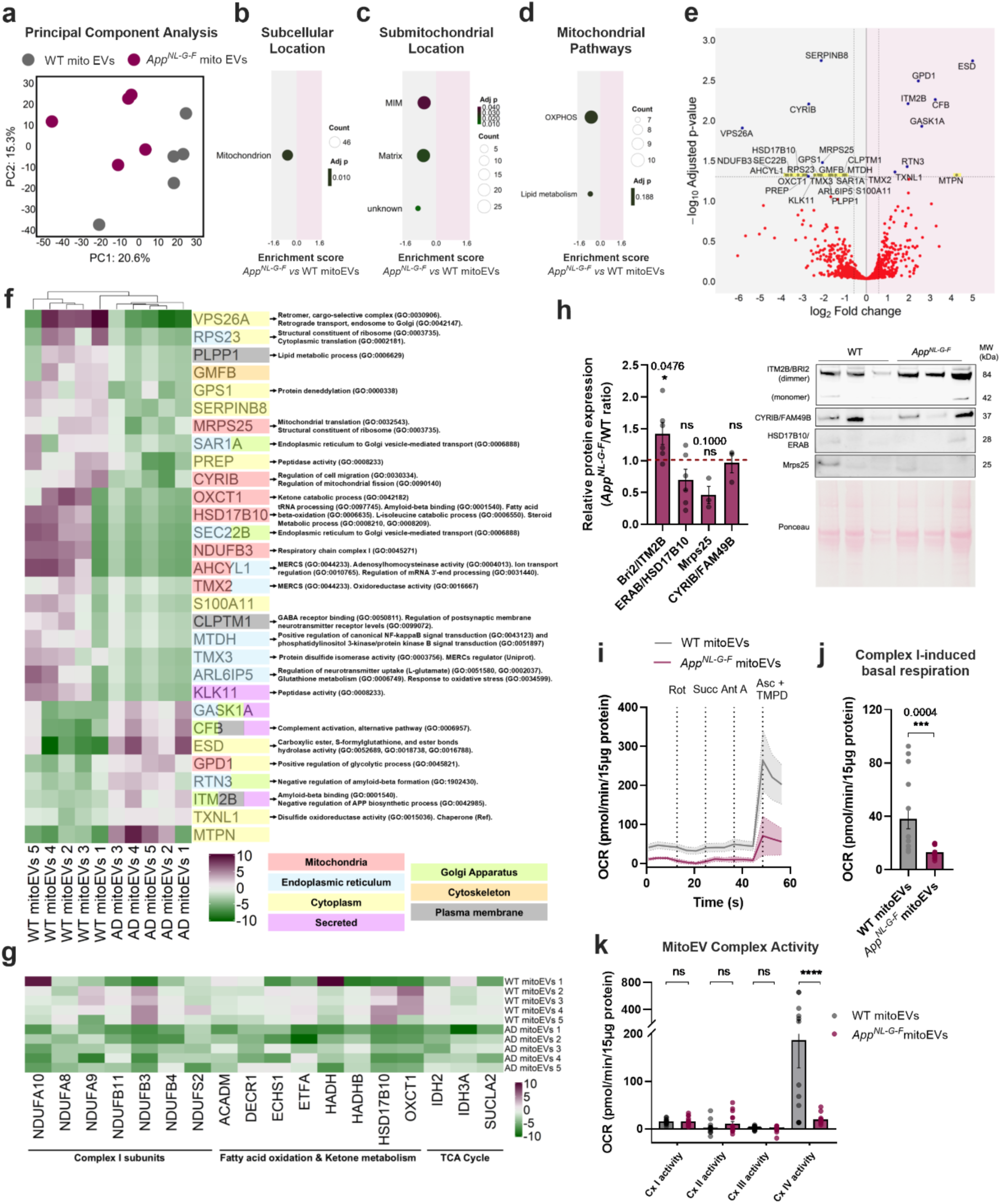
*App^NL-G-F^*astrocyte-derived mitoEVs exhibit deficient mitochondrial protein cargo and function. **(a)** PCA plot of WT and *App^NL-G-F^* mitoEVs. **(b-d)** Dotplots of GSEA enrichment scores comparing *App^NL-G-F^ versus* WT mitoEVs for **(b)** subcellular location, **(c)** submitochondrial location, or **(d)** mitochondrial pathway terms. Terms downregulated in AD relative to WT appear on the left. Dot size and color indicate the number of associated proteins and GSEA adjusted *p*-value, respectively. **(e)** Volcano plot comparing *App^NL-G-F^* and WT mitoEV proteomes. Proteins enriched in *App^NL-G-F^* relative to WT appear on the right, while downregulated proteins appear on the left. Y- and x-axes show the -log_10_ adjusted p-value and log_2_ fold change, respectively; dashed lines indicate significance thresholds. Yellow highlights proteins not detected in one of the genotypes. **(f-g)** Heatmaps representing **(f)** significantly altered proteins and **(g)** representative proteins associated with mitochondrial pathway subterms in WT and *App^NL-G-F^*/AD mitoEV samples. The color gradient encodes the row-centered intensities of each protein across all samples. In **(f)**, the colored backgrounds of protein labels indicate known subcellular localizations, and labels on the right denote selected GO terms for each protein. **(h)** Representative Western blots and quantification of selected differentially expressed proteins in mitoEVs. Data are shown as the *App^NL-G-F^* / WT ratio of relative protein expression (n = 3–6). **(i-k)** OCR was measured in mitoEVs using the Seahorse assay. **(i)** Representative OCR traces following sequential injections of rotenone (2 µM), succinate (10 mM), antimycin A (4 µM), and ascorbate (1 mM)/TMPD (100 mM). **(j)** Complex I–driven OCR. **(k)** Complex I–IV activities (n = 9). Data presented as mean ± SEM, and statistical significance was determined using 2-way ANOVA. * p ≤ 0.05, *** p ≤ 0.001, **** p ≤ 0.0001, ns. non-significant.

To further resolve the molecular basis of these changes, we examined differentially abundant proteins using volcano plot analysis from the hybrid dataset (**Fig 5e**). This analysis showed 16 proteins absent in *App^NL-G-F^* but present in WT mitoEVs (e.g., Ndufb3, Hsd17b10/Erab, Oxct1) and one *App^NL-G-F^* mitoEV-specific protein (Mtpn). Among those shared between *App^NL-G-F^* and WT mitoEVs, seven were significantly enriched in *App^NL-G-F^* mitoEVs (e.g., Gpd1, Itm2b/Bri2, Rtn3) while six were significantly depleted (e.g., *Vps26a, Mrps25,* Cyrib/Fam49b). GO functional annotation of these differentially expressed proteins highlighted enrichment of factors linked to negative regulation of APP-related processes (e.g., Itm2b/Bri2)[41], while depleted proteins were predominantly associated with MIM and matrix compartments, mitochondria–ER contact sites (MERCS; e.g., Ahcyl1), mitochondrial ribosomal components and mitoEV/MDV biogenesis (e.g., Vps26a) (**Fig 5e, f**). In addition, proteins involved in oxidative metabolism (mainly involved in Complex I), lipid and fatty acid metabolism, redox homeostasis, and mitochondrial gene expression were underrepresented or absent in *App^NL-G-F^*mitoEVs (**Fig 5g, Supplementary Fig S4f**). Immunoblotting of selected differentially represented proteins confirmed an upregulation of Itm2b/Bri2 (p = 0.0476) and trends toward downregulation of Hsd17b10/Erab (p = 0.3636) and the mitochondrial ribosome protein Mrps25 (p = 0.1), reflecting high inter-sample variability (**Fig 5h**).

Recent observations in yeast suggest that mitoEVs retain mitochondrial membrane potential and functional ATP synthase^7^. To assess whether the reduced mitochondrial inner membrane content and depletion of OXPHOS proteins observed in *App^NL-G-F^* astrocytic mitoEVs impact their bioenergetic capacity, we established a Seahorse-based assay to measure uncoupled respiration and individual respiratory complex activity in isolated mitoEVs. Respiration was driven by pyruvate and malate to support Complex I activity, followed by sequential application of specific inhibitors and substrates to examine the individual complexes. While Complex I–driven basal respiration showed a marked reduction in *App^NL-G-F^* mitoEVs (p = 0.0004), Complex I activity was preserved (**Fig 5i, j**). In contrast, Complex IV activity was strongly reduced in *App^NL-G-F^*mitoEVs (p < 0.0001). Notably, mitoEV respiration was insensitive to antimycin A, indicating a lack of functional Complex III activity in astrocyte-derived mitoEVs. **(Fig 5i, k)**.

Taken together, these findings demonstrate that *App^NL-G-F^* astrocyte-derived mitoEVs are selectively depleted of key mitochondrial structural and bioenergetic components, resulting in impaired respiratory capacity, particularly at Complex IV.

### *App* knock-in-derived mitoEVs fail to confer metabolic support to neurons

Disease-associated alterations in cargo and bioenergetic capacity in WT and *App^NL-G-F^* astrocyte-derived mitoEVs prompted us to evaluate their ability to support neuronal metabolic function. To determine an appropriate treatment dose, neuronal metabolic viability was assessed 24 h after mitoEV exposure using the CellTiter-Glo assay, which measures intracellular ATP levels. MitoEV concentrations ranging from 5.6 to 96 pg per neuron were tested, with higher doses resulting in reduced ATP levels (**Fig 6a, b, supplementary Fig S5a, b**). Based on these results a concentration of 12 pg mitoEVs per neuron was selected for subsequent experiments.

**Figure 6:**
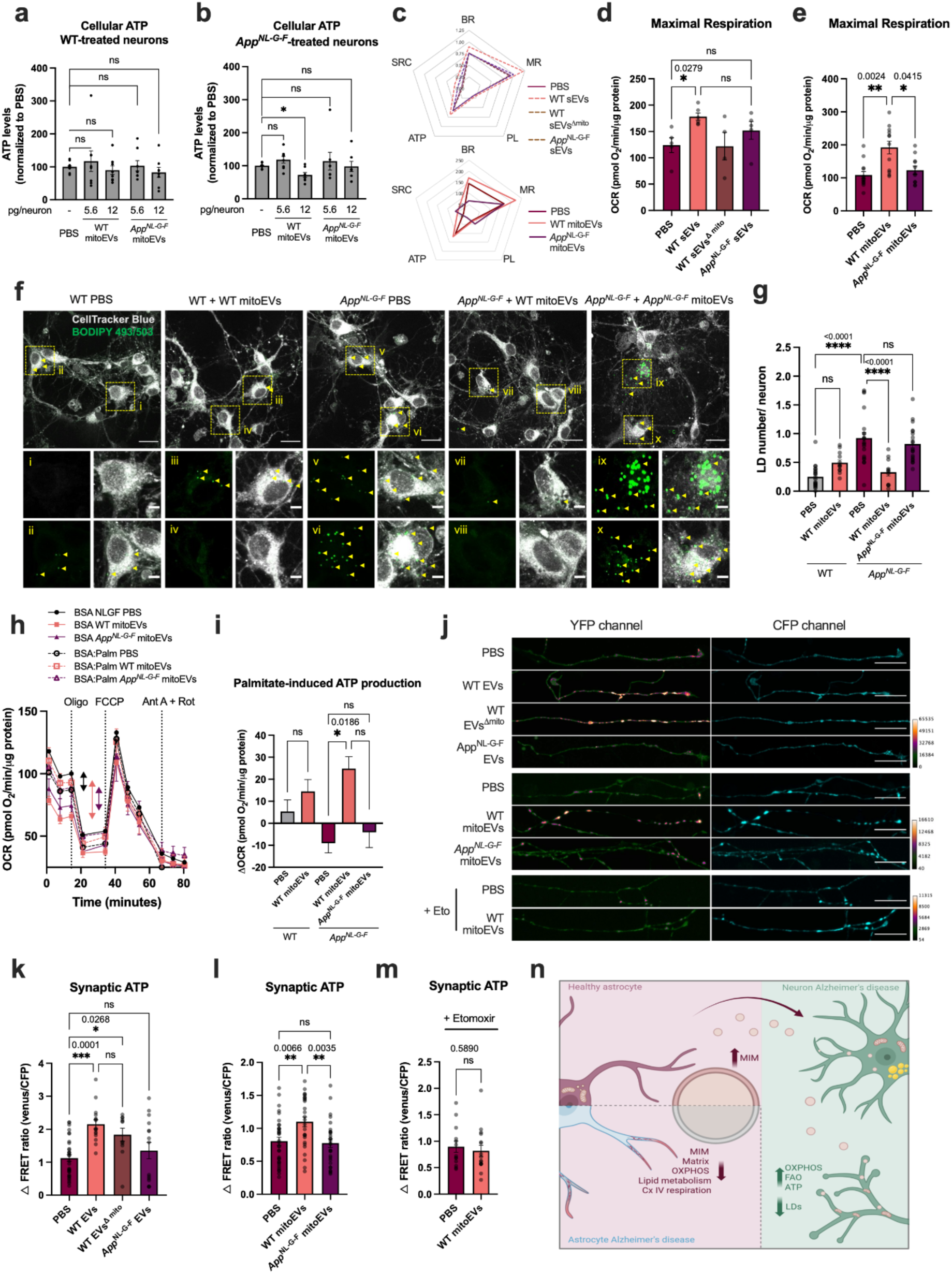
MitoEVs affect bioenergetics and lipid handling in AD neurons. Neurons were treated, when indicated, with astrocyte-derived EVs (0.5 ng/neuron, 6 h) or mitoEVs (5.6 or 12 pg/neuron, 24 h) (n = 5-6). **(a, b)** Cellular ATP levels were quantified using Cell Titer-Glo in **(a)** WT and **(b)** *App^NL-G-F^* treated neurons. **(c-e)** OCR was evaluated using the Seahorse assay (n = 5-7). Functional mitoEVs were depleted from the EV pool (EVs^△mito^) by treating the EVs with the mitochondria uncoupler FCCP (2 µM, 30 min). **(c)** Spider charts depict the distribution of Seahorse parameters across conditions, including basal respiration (BR), maximal respiration (MR), proton leak (PL), ATP-linked respiration (ATP), and spare respiratory capacity (SRC). **(d, e)** Bar graphs represent the maximal respiration of treated neurons. **(f, g)** Lipid droplet number was quantified by confocal microscopy using BODIPY 493/503 staining. Neuronal surface area was determined using CellTracker Blue (n = 21-30 from 3 independent cultures). Scale bars = 20 µm; 5 µm in insets. **(h, i)** OCR was measured using the Seahorse assay in neurons cultured in low-glucose medium and stimulated with palmitate (1 mM, 4 h). Basal and maximal respiration were quantified based on palmitate-induced increases in OCR. **(j-m)** Synaptic ATP levels were quantified using the FRET-based ATP sensor SypI-ATeam1.03. FRET ratio (Venus/CFP) was calculated in neurons at rest (n = 21-30 from 3 independent cultures) and in the presence of etomoxir (Eto) when indicated on the graph. Scale bar = 20 µm. **(n)** Schematic representation of the hypothesis. Figure made using BioRender.com. Data presented as mean ± SEM, and statistical significance was determined using ANOVA Šídák’s multiple comparisons test or Kruskal-Wallis test. * p ≤ 0.05, ** p ≤ 0.01, *** p ≤ 0.001, **** p ≤ 0.0001, ns. non-significant.

EVs have previously been shown to modulate mitochondrial function[42], however, the contribution of specific EV subpopulations remains unclear. We first treated *App^NL-G-F^* neurons with total EVs derived from WT and *App^NL-G-F^* astrocytes. To assess the specific contribution of mitoEVs, functional mitochondrial activity within WT EV preparations was selectively disrupted by FCCP treatment (WT EVs^Δmito^). WT EVs significantly increased both mitochondrial maximal respiration (p = 0.0279) and spare respiratory capacity, whereas neither *App^NL-G-F^* EVs nor WT EVs^Δmito^, affected the respiration (**Fig 6c, d, Supplementary Fig S5c-e**). These findings indicate that functional mitoEVs are required for the bioenergetic effects of astrocyte-derived EVs on neurons. Therefore, we treated neurons with purified WT and *App^NL-G-F^* mitoEVs. WT mitoEVs significantly increased neuronal maximal respiration (p = 0.0173), whereas *App^NL-G-F^* mitoEVs had no effect (**Fig 6c, e, Supplementary Fig S5f-h**). These findings indicate that functional mitoEVs are required for the bioenergetic effects of astrocyte-derived EVs on neurons and that disease-associated alterations in *App^NL-G-F^* mitoEVs impair their ability to support neuronal mitochondrial function.

### MitoEVs promote lipid droplet mobilization to fuel fatty acid oxidation in App neurons

Lipid droplet (LD) accumulation has been reported in both AD mouse models and patient brains, predominantly in glial cells[43, 44]. Although the use of lipids as an energy source by neurons has been debated, previous work from our laboratory using a similar *App* knock-in model demonstrated that neurons can also accumulate LDs [45]. Our proteomic data showed downregulation and/or absence of key FAO-related enzymes in *App^NL-G-F^*-derived mitoEVs (**Fig 5f, g; Supplementary Fig S4h, i**), prompting us to investigate whether mitoEVs modulate neuronal lipid metabolism. BODIPY staining revealed a marked increase in LD number in *App^NL-G-F^* neurons compared with WT controls (p < 0.0001), which was restored to basal levels following treatment with WT mitoEVs (p < 0.0001) but not with *App^NL-G-F^* mitoEVs (**Fig 6f, g, Supplementary Fig S5i**). Because LD accumulation is commonly associated with impaired long-chain fatty acid utilization and FAO [46], we next assessed neuronal FAO capacity using Seahorse analysis with palmitate as the primary substrate. *App^NL-G-F^* neurons displayed a profound inability to utilize fatty acids compared with WT neurons (**Fig 6h, i**), consistent with increased LD accumulation. Notably, WT mitoEV treatment enabled *App^NL-G-F^* neurons to increase both basal and ATP-linked respiration in the presence of palmitate, an effect not observed with *App^NL-G-F^*mitoEVs or control treatments (**Fig 6h, i**).

Recent evidence suggests that mitochondria can utilize fatty acids mobilized from axonal LDs to support local ATP production[46, 47]. To test whether mitoEV-mediated metabolic rewiring translates into improved synaptic energy availability, we measured synaptic ATP levels. Under identical treatment conditions as reported in **Fig 6d**. We observed that neurons treated with WT astrocyte-derived EVs exhibited an approximately two-fold increase in synaptic ATP levels (p = 0.0002), whereas EVs derived from *App^NL-G-F^* astrocytes had no significant effect (**Fig 6j, k**). EVs^Δmito^ retained only a modest ability to elevate synaptic ATP (p = 0.04), suggesting mitochondrial activity is an important, but not exclusive, contributor to the increased synaptic ATP (**Fig 6j, k**). Consistent with this, treatment with purified WT – but not *App^NL-G-F^* – mitoEVs resulted in increased synaptic ATP levels (**Fig 6j, l**). This effect was lost in the presence of etomoxir, an irreversible inhibitor of CPT1, which blocks the long-chain fatty acid import into the mitochondria (**Fig 6m**). These findings show that the mitoEV-mediated increase in synaptic ATP requires the import and oxidation of long-chain fatty acids.

Taken together, these findings demonstrate that WT astrocytic mitoEVs support neuronal bioenergetics by promoting LD mobilization, FAO, and mitochondrial respiratory capacity, with associated improvements in synaptic energy availability (**Fig 6n**). In contrast, *App^NL-G-F^*-derived mitoEVs fail to engage these metabolic pathways, underscoring a disease-associated impairment in astrocyte-to-neuron metabolic support in AD.

## DISCUSSION

Mitochondrial impairment is increasingly recognized as an early feature of AD [4, 45], yet whether changes in mitochondrial dynamics drive bioenergetic failure or reflect compensatory adaptations remains unclear. Here, we identify a spatially restricted bioenergetic deficit in *App^NL-G-F^* neurons characterized by presynaptic mitochondrial loss and reduced synaptic ATP despite preserved global ATP levels, while astrocytes exhibit a more dynamic but metabolically preserved mitochondrial network. Importantly, we show that astrocyte-to-neuron mitochondrial transfer directly modulates neuronal bioenergetics and provide the first comprehensive structural, proteomic, and functional characterization of astrocyte-derived mitoEVs. These vesicles are enriched in OXPHOS, lipid metabolism, and detoxification pathways and retain measurable respiratory capacity. In contrast, *App^NL-G-F^*-derived mitoEVs are selectively depleted of these bioenergetic components and fail to promote neuronal lipid droplet mobilization, fatty acid utilization, and OXPHOS-dependent respiration. Together, our findings identify mitoEV-mediated mitochondrial transfer as a previously unrecognized mechanism of neuron–glia metabolic coupling that is compromised in early AD and reveal a coordinated role for OXPHOS and FAO pathways in supporting synaptic energy homeostasis.

A network-based analysis modeling neuronal vulnerability in AD identified axon terminals as the most susceptible module, closely associated with microtubule-based transport and synaptic plasticity [48]. Mitochondria play a central role in this module, as a substantial proportion of neuronal ATP is consumed to sustain axonal transport and maintain resting membrane potential in both neurons and glia [49]. Although mitochondria are highly dynamic organelles whose fission, fusion, and trafficking enable their redistribution within polarized cells [50], not all synapses contain resident mitochondria, and how mitochondrial supply matches rapidly shifting local energy demands remains incompletely understood. One potential mechanism is horizontal mitochondrial transfer from neighboring cells, such as astrocytes. Here we show that *App^NL-G-F^* astrocytes exhibit increased mitochondrial trafficking and remodeling, and – when co-cultured with *App^NL-G-F^* neurons – upregulate genes associated with mitochondrial transport and biogenesis. Similarly, previous studies have shown that the astrocytic mitochondrial network adapts to neuronal activity in the vicinity of the synapse [51]. In the context of AD, such adaptive remodeling may prime astrocytes for increased mitochondrial turnover or redistribution in response to the heightened presynaptic excitatory and inhibitory activity observed at early disease stages, including in *App^NL-G-F^* mice [52].

Emerging evidence suggests that mitochondrial transfer from glial cells to neurons supports neuronal survival under pathological conditions, with glia-derived mitochondria reported to improve neuronal function by replenishing damaged mitochondrial pools, mainly in models of ischemic stroke and Parkinson’s disease [9, 11, 53]. A couple of studies have shown similar results after intravenous injection of isolated mitochondria in rodent models of AD, with increased neuronal and mitochondrial function [54, 55]. Although the precise mechanisms underlying horizontal mitochondrial transfer remain incompletely defined, TNTs and EVs are considered the primary pathways [12, 36, 56, 57]. TNT-mediated mitochondrial exchange has been described between neurons and glial cells, including under α-synuclein toxicity [12]; however, we did not detect TNT formation in our co-culture system. Instead, vesicle-mediated transfer appears to be the dominant route for transfer. While we cannot formally exclude a minor contribution of TNTs – given the technical challenges in visualizing these structures in complex cellular networks – our imaging analyses consistently revealed punctate vesicular mitochondrial structures rather than filamentous transfer events. Notably, recent work has shown that cells can engage both TNT- and EVs-mediated mechanisms [56], suggesting that multiple pathways may coexist.

In our experimental paradigm, mitochondrial transfer efficiency was significantly reduced in *App^NL-G-F^* cultures in both directions. Despite previous reports showing that stress and disease conditions enhance mitoEV release, our data suggest that this reduction was not due to altered mitoEV release [14]. Rather, impaired transfer likely reflects qualitative alterations in mitoEV composition, including changes in protein cargo, membrane properties, or uptake efficiency by recipient cells. One possibility is that disease-associated factors at the vesicle surface modulate recognition and uptake. Aβ has been detected in both EV [21] and mitochondrial membranes[58], potentially altering vesicle bioactivity or cellular uptake. Likewise, surface-associated enzymes such as MAO-B [22], which influence synaptic redox signaling, may affect mitoEV–neuron interactions. Although speculative, these mechanisms suggest that altered mitoEV surface composition could contribute to defective mitochondrial transfer in AD.

The exact nature of mitoEVs remains to be fully defined. They may derive from MDVs, which bud directly from mitochondrial membranes as part of mitochondrial quality control pathways [17] [59, 60]. Rather than being targeted for degradation, a subset of MDVs might be rerouted to the endosomal pathway and released as mitoEVs [16, 17]. Structural similarities between MDVs and mitoEVs – including their small size, absence of cristae, and lack of proteins such as TOM20 and Mfn2 – support this hypothesis [16, 61]. Here, we provide an in-depth proteomic and functional characterization of astrocytic mitoEVs. WT mitoEVs are protein-dense vesicles enriched in MIM and matrix components linked to OXPHOS, the TCA cycle, amino acid and lipid metabolism. Consistent with this, recent studies suggest that mitoEVs retain mitochondrial membrane potential and the ability to produce ATP [10, 16, 18]. In addition, they contain proteins involved in ROS detoxification and glutathione metabolism, suggesting that mitoEVs may provide antioxidant capacity capable of buffering redox imbalance during neurodegenerative stress [62]. In agreement, inter-organ mitochondrial vesicle transfer was shown to enhance antioxidant resilience in recipient cells [10], supporting a conserved redox and protective function. The presence of non-mitochondrial proteins from the plasma membrane, ER, and cytosol may reflect biogenesis at mitochondrial contact sites, including MERCs and peroxisomal interfaces. However, partial contamination from sEV fractions represents a technical limitation of the isolation approach and is likely to contribute to this profile.

In contrast, *App^NL-G-F^* astrocytic mitoEVs display marked downregulation of MIM and matrix proteins, including respiratory chain subunits and enzymes required for lipid and fatty acid metabolism. This qualitative shift is accompanied by functional impairment, particularly reduced Complex IV activity, the respiratory complex most consistently affected in AD brains [63]. Notably, our previous work demonstrated increased Complex IV respiration in pre-symptomatic *App^NL-G-F^* brains [4], raising the possibility that mitoEV secretion may contribute to mitochondrial quality control by selectively exporting dysfunctional respiratory components. Additionally, several FAO-related proteins reduced/absent in *App^NL-G-F^* mitoEVs, such as Hsd17b10, are also decreased in CSF from AD patients [64], underscoring the clinical relevance of these cargo alterations. Notably, Hsd17b10 (also known as ERAB/ABAD) interacts with Aβ in the mitochondrial matrix and has been implicated in mitochondrial dysfunction in AD [3], linking mitoEV cargo alterations to pathways associated with Aβ-induced mitochondrial toxicity. *App^NL-G-F^* mitoEVs further exhibit increased levels of Itm2b/Bri2, a protein known to interact with APP and to act as an inhibitor of Aβ oligomerization [65, 66]. Bri2 is increased in amyloid plaques in early stages of AD [67] and loss of Bri2 was linked to synaptic and memory dysfunction in mouse models of dementia [68]. The increased shuttling of Bri2 into the mitoEVs may be an attempt of the astrocytes to protect neurons and synapses of Aβ oligomerization, or could indicate altered cargo-selection mechanisms in AD.

Our proteomic analysis revealed the absence of key mitochondrial fusion proteins (e.g., Mfn1, Mfn2, OPA1) in astrocyte-derived mitoEVs, which may explain their limited integration into the recipient mitochondrial network. However, integration may not be required for functional impact, as mitochondrial components within EVs can modulate cellular metabolism by supplying bioenergetic or redox-active molecules that enhance OXPHOS and antioxidant defense [10, 69]. Consistently, metabolic profiling of *App^NL-G-F^* brain-derived EVs has revealed reductions in metabolites linked to neurotransmission and redox balance (e.g., GABA, glutamic acid, glutathione) [70], suggesting that altered vesicle composition may directly influence neuronal metabolic support. We further speculate that non-integrated mitoEVs may contribute to mitochondrial quality control during neuron-to-astrocyte transfer, as previously described in AD and Parkinson’s disease models [39, 71].

Previous studies examining mitochondrial transfer in the brain, including in tau- and α-synuclein models, have largely emphasized the restoration of OXPHOS-related processes [12, 53]. Consistently, we show that WT mitoEVs restore maximal respiration in App^NL-G-F^ neurons, indicating that these vesicles can compensate for intrinsic mitochondrial deficits. Notably, our proteomic analysis revealed that a substantial fraction of mitoEV cargo is linked not only to OXPHOS and carbohydrate metabolism, but also to lipid metabolic pathways, including FAO, many of which are reduced or absent in *App^NL-G-F^* mitoEVs. Although glucose has been widely regarded as the primary fuel source for neurons [72], increasing evidence indicates that neurons can engage FAO under conditions of metabolic stress or elevated activity to support synaptic function and memory formation [46, 47] [73–75]. While LD accumulation in AD has been primarily associated with glial pathology [76, 77], we have shown that AD neurons also accumulate LD [45]. Here, we demonstrate that *App^NL-G-F^* neurons fail to engage FAO under our experimental conditions, suggesting impaired mobilization of LD-derived substrates to mitochondria. Importantly, WT – but not *App^NL-G-F^* – mitoEVs restore LD mobilization and FAO capacity, increasing both basal and ATP-linked respiration. This metabolic rewiring is especially relevant at the level of the synapse, where fatty acid utilization occurs in an activity-dependent manner, supporting local ATP production during heightened energetic demand [46, 47, 75]. In line with this, WT mitoEV treatment significantly increased synaptic ATP levels in a manner dependent on mitochondrial fatty acid import, indicating that the mitoEV-mediated enhancement of synaptic ATP is likely mediated by coordinated OXPHOS and FAO pathways. Together, these findings uncover a previously unrecognized role for astrocytic mitoEVs in reinforcing neuronal LD mobilization and fatty acid oxidation, linking intercellular mitochondrial transfer to synaptic energy supply.

## CONCLUSION

Our findings establish mitoEV-mediated mitochondrial transfer as a previously unrecognized mechanism by which astrocytes regulate several neuronal bioenergetic pathways, including mitochondrial oxidative phosphorylation, lipid handling, and subsequent synaptic energy availability, expanding the concept of neuron-astrocyte metabolic coupling. Importantly, we demonstrate that this mechanism is impaired in early AD, linking mitochondrial communication between neurons and glia to early synaptic vulnerability.

This study deepens our understanding of mitochondrial transfer in the brain, but at the same time raises important questions about the specificity and regulation of mitoEV cargo selection. Determining whether these vesicles mirror astrocytic metabolic stress, or whether their contents are meticulously selected to support neuronal bioenergetics will be essential to understanding their role in physiological circumstances and the potential to harness them for therapeutic or biomarker purposes. Additionally, further *in vivo* studies will be necessary to validate this mechanism and define its regulation in AD. Notably, the comprehensive proteomic profiling of astrocytic mitoEVs presented here offers a valuable resource to guide the design of future *in vivo* strategies to study mitoEV-mediated mitochondrial transfer.

## Declarations

### Consent for publication

Not applicable.

## Availability of data and materials

The mass spectrometry proteomics data have been deposited to the ProteomeXchange Consortium via the PRIDE partner repository [78] with the dataset identifier PXD074664. The proteomic analysis and mitochondrial 4D network imaging analysis code are available in a GitHub repository and will be made publicly available upon publication. Other data published in this study are available from the corresponding author upon request.

## Authors’ contributions

L.N. conceptualized the study; L.V., R.G., F.P., M.J.P., S.G., Z.C., K.D., D.K.M. and D.R.M., performed the experiments; D.K.M., O.P.B.W. and D.R.M. provided technical assistance in EV isolation; M.M and J.B.P. provided support in bioinformatic analysis; L.V., R.G., J.A.G., F.P., M.J.P., S.G., K.D. Z.C. and L.N. analyzed data; L.V., M.A. and L.N. coordinated the project; M.A. and L.N. provided funding; L.V., J.A.G., and L.N. wrote the manuscript with contributions from all authors.

## Acknowledgements

We thank Takaomi Saido and Takashi Saito at RIKEN Center for Brain Science (Japan) for providing App knock-in mice. We also thank the Beta Cell in-vivo Imaging/Extracellular Flux Analysis core facility (supported by the SRP Diabetes) for providing support with Seahorse experiments, the Biomedicum Imaging Core Facility (BIC) staff for their input with imaging analysis, the EM facility in Huddinge Hospital (EMil) and the cryo-EM Swedish National Facility funded by the Knut and Alice Wallenberg, Family Erling Persson and Kempe Foundations, SciLifeLab, Stockholm University and Umeå University, for valuable help with EM experiments. We are grateful to Professor Susanne Gabrielsson and Dr. Loïc Steiner from Karolinska Institutet for providing access to the NTA and TFF systems. We thank Dr. Daniel Gitler from the Zlotowski Center for Neuroscience (Israel) by kindly providing aTeam FRET sensors, and Dr. Simone Tambaro from Karolinska Institutet for providing the ITM2B/Bri2 antibody. Technical University of Denmark and Biogenity has contributed by performing the proteomics sample analysis.

