## Supplementary data for "Astrocyte-neuron mitochondrial transfer via mitoEVs supports neuronal energy metabolism and is impaired in early Alzheimer’s disease"

### Supplementary files

**Supplementary Methods**

**Mitochondrial 4D network imaging analysis in astrocytes**

Time-lapse image stacks acquired by spinning-disk confocal microscopy were processed using a custom computational pipeline implemented in Python (Jupyter Notebook, Anaconda distribution; available in a GitHub repository https://github.com/jesusalarcongil/AstrocyteNeuronMitoEVs upon request). First, the original time lapses (nd2) were transformed to independent ".tiff" files per frame thanks to the bfconvert (bftools) function from the consortium of Open Microscopy Environment[1]. Afterward, these independent files were deconvoluted with the DeconWolf tool[2], using the following microscope settings: numerical aperture: 1.42, refractive index: 1.515, emission wave: 583, lateral resolution: 111 nm, and axial resolution: 200 nm, and twenty iterations per image. The deconvolved images were thresholded to remove background signal and the mitochondrial network was segmented using MitoGraph[3].

The MitoGraph output for each time frame from all time-lapse recordings was then used as input for MitoTNT[4], which was run in Jupyter Notebook according to the developers' instructions. Briefly, segmented mitochondrial networks from frames 0–30 (frame interval: 2 s) were prepared with a node gap size of 20 to enable mitochondrial network tracking across time using a gap-closing function (min_track_size: 2; max_gap_size: 1). These tracked mitochondrial networks were used to calculate mitochondrial fusion and fission events as well as mitochondrial segment diffusivity.

For the analysis of mitochondrial morphology and network architecture, the output generated for the first temporal frame of each time-lapse was used as input for the scripts available at https://github.com/Hill-Lab/MitoGraph-Contrib-RScripts, implemented in RStudio (R) on a Windows system. In this framework, segmented mitochondria are represented as a graph composed of nodes (branch points or termini) connected by edges (mitochondrial segments). From this skeletonized representation, quantitative descriptors of mitochondrial morphology and network topology were extracted, including total nodes, total edges, total network length, mitochondrial volume estimates, average edge length and width, number of free ends, three-

and four-way junctions, and average node degree, providing quantitative measures of mitochondrial size, branching, and connectivity.

**Mass spectrometry (MS) sample preparation and analysis**

To eliminate residual iodixanol, isolated mitoEVs were run briefly into a 12% Bis-Tris gel (Invitrogen, #NP0341BOX) in NuPAGE MES SDS running buffer (ThermoFisher, #NP0002) until all proteins focused into a single 1-cm band. Gels were washed three times in dd$H_2O$ for 15 min each and visualized by staining with EZBlue™ Gel Staining Reagent (Sigma, #G1041). Gel bands were excised into small pieces using a scalpel. The gel pieces were destained for 30 minutes in a destaining buffer (50% acetonitrile in 50 mM ammonium bicarbonate), with occasional vortexing to ensure complete destaining. Once the color was fully removed, the gel pieces were dehydrated by adding 100% acetonitrile and vortexing. The solvent was discarded, and the gel pieces were rehydrated in a reduction solution (10 mM TCEP, 40 mM CAA, and 80 mM ammonium bicarbonate) and incubated with occasional vortexing for 30 minutes. 500mM Acetonitrile was added to dehydrate the gel pieces and excess liquid was discarded. Any residual solvent was removed by evaporation in a sterile bench. Trypsin solution (25ng/µL trypsin (MS grade, Sigma) in 10mM AMBIC and 10% Acetonitrile) was added and samples were incubated on ice for 20 min. Trypsin solutions were then discarded and 100mM AMBIC was added to the samples. Samples were then incubated at 37 °C overnight. The following day, extraction buffer (5% Formic Acid in 50% Acetonitrile) was added to the samples, which were then incubated at 37 °C for 30 min. Supernatants containing the peptides were transferred to clean Lo-Bind Eppendorf tubes. Extractions were repeated and the pooled supernatants were concentrated in an Eppendorf Speedvac. The concentrated peptides were reconstituted in 30 µL of solution A* (2% acetonitrile, 1% trifluoroacetic acid) for analysis. The peptides were then centrifuged at 18,000 × g for 10 minutes to remove any residual particulate matter. The supernatant was carefully transferred and analyzed using a Nanodrop to determine peptide concentration.

Peptides were loaded onto a 2 cm C18 trap column (ThermoFisher 164946), connected in-line to a 15cm C18 reverse-phase analytical column (Thermo EasySpray ES904) using 100% Buffer A (0.1% Formic acid in water) at 750bar, using the Thermo EasyLC 1200 HPLC system, and the column oven operating at 30 °C. Peptides were eluted over a 70 minute gradient ranging from 10% to 60% of Buffer B (80% acetonitrile, 0.1% formic acid) at 250 nl/min, and the Orbitrap Exploris instrument (Thermo Fisher Scientific) was run in data-independent acquisition (DIA) mode with FAIMS ProTM Interface (ThermoFisher Scientific) with CV of -45 V. Full MS spectra were collected at a resolution of 120,000, with an AGC target of 300% or maximum injection time set to 'auto' and a scan range of 400–1000 m/z. The MS2 spectra were obtained in DIA mode in the orbitrap operating at a resolution of 60.000, with an AGC target 1000% or maximum injection time set to 'auto', a normalized HCD collision energy of 32. The isolation window was set to 6 m/z with a 1 m/z overlap and window placement on. Each DIA experiment covered a range of 200 m/z resulting in three DIA experiments (400-600 m/z, 600-800 m/z and 800-1000 m/z). Between the DIA experiments a full MS scan was performed. MS performance was verified for consistency by running complex cell lysate quality control standards, and chromatography was monitored to check for reproducibility. Technical University of Denmark, through Biogenity, has contributed by performing the proteomics sample analysis.

**LC-MS Proteomic Data Analysis**

The raw files were analyzed using SpectronautTM (version 19.1), spectra were matched against the *Mus musculus* database (Uniprot taxonomi 10090). The proteomic data analysis was performed in R 4.4.2 using various Bioconductor 3.20 packages (e.g., DEP[5] and clusterProfiler[6]) and built-in functions. The code, original data, input files, and raw outputs from this analysis are available in a GitHub repository (https://github.com/jesusalarcongil/AstrocyteNeuronMitoEVs) and can be accessed upon request. The identified protein intensities provided by Biogenity were imported into R and restructured to isolate the experimental group metadata and the protein intensities, together with their symbols and UniProt mouse identifiers. Additionally, to support the downstream analysis, we generated three custom term-annotation databases. One of them included terms on the protein's subcellular localization, integrated from the Human Protein Atlas[7] , UniProt[8], and Hein *et al.* 2025[9]. The other two custom term-annotation databases included terms related to the protein's submitochondrial localization and mitochondrial pathways, which were imported from MitoCarta 3.0[10] . For building these custom term-annotation databases, and for other downstream analyses, mouse and human protein identifiers (Symbol, UniProt ID, and Entrez ID) were mapped using the mapIds function, the homologene package, and manual curation of unmapped IDs and homologues.

First, we profiled the proteins identified in WT-derived mitoEVs (all proteins identified in the analysis, excluding the one identified only in *App*$^{NL\text{-}G\text{-}F}$-derived mitoEVs). We performed Over-Representation Analysis (ORA) of the WT-derived mitoEVs protein list against different annotation databases (KEGG[11], GO Biological Process (GO BP), GO Cellular Component (GO CC), GO Molecular Function (GO MF)[12], custom Subcellular Location annotations, custom Submitochondrial Location annotations, and custom Mitochondrial Pathway annotations), using mouse as the organism and the proteins in the organism or in each annotation database as the universe for the analysis. To perform these analyses, we used the functions *enrichKEGG*, *enrichGO*, or *enricher,* with an annotation size filter ranging from 3 to 30,000. Additionally, we adjusted the resulting p-values using the False Discovery Rate (FDR). Since we identified a considerable number of overrepresented terms in the KEGG, GO BP, GO CC, and GO MF ORA, we clustered these terms using an adjusted p-value threshold < 0.125, a minimum of 2 or 3 genes per term, and a minimum cluster size ranging from 5 to 40 terms. We performed this clustering using the *simplifyEnrichment* package for GO annotations and the gene overlap coefficient for KEGG annotations. Each cluster was labelled with four manually selected keywords among them the terms included in each cluster. All ORA results were plotted as dot plots.

Subsequently, the identified proteins were divided into two groups based on their missingness patterns for a hybrid differential enrichment analysis comparing *App*$^{NL\text{-}G\text{-}F}$- and WT-derived mitoEVs (*App*$^{NL-G-F}$- *versus* WT-derived mitoEVs). Proteins detected at least once in each group were analyzed using the *DEP* package. Briefly, the missing protein intensities – presumed to be primarily missing not at random due to low protein concentrations in EV samples – were imputed using random values drawn from a Gaussian distribution with a mean at 0.05 quartile for each sample. Thereafter, a Variance Stabilization Normalization (VSN) matrix was computed to normalize the data. Following that, a differential enrichment analysis was performed using protein-specific linear models and empirical Bayes statistics, with false discovery rate-adjusted p-values, log2 fold changes, and centered log2 intensities. Conversely, proteins not detected in any sample within a group were imputed using the observed limit of detection for each sample, as their presence in that group could not be verified. These data were then normalized using the previously calculated VSN matrix. With that, log2 fold changes and centered log2 intensities were calculated. A presence/absence contingency table was generated from the original protein intensity data, and with it, a Fisher's exact test with false discovery rate correction was performed to obtain the adjusted p-values. Combining the results, we plotted the first two principal components from Principal Component Analysis (PCA) to explore potential differences between groups and samples. A correlation plot was also generated to examine the differences between groups. Then, a Volcano Plot showing the -log10 of the adjusted p-value versus the log2 fold change for each protein in the *App*$^{NL-G-F}$- versus WT-derived mitoEVs comparison was generated. A threshold of -log10(0.05) for the adjusted p-value and a threshold of log2(1.5) for the fold change were used to consider a protein significantly enriched or depleted in the *App*$^{NL-G-F}$- versus WT-derived mitoEVs. Furthermore, heatmaps of log2-centered intensities for selected proteins (i.e., those significantly enriched or depleted in *App*$^{NL-G-F}$-derived mitoEVs) were generated across samples. The subcellular location and GO terms plotted in the heatmap were manually retrieved from UniProt. Doughnut charts were also generated to show the relative percentages of VSN-normalized protein abundances associated with terms from the three custom annotation databases in WT astroglial mitoEVs. Each segment represents the sum of VSN-normalized protein abundances for a given term, expressed as a percentage of the total VSN-normalized abundances of proteins with assigned terms in WT astroglial mitoEVs. The color legend maps each term to its corresponding segment. Labels within each segment indicate the percentage contribution of that term. Overlay maps of selected Reactome pathways were also generated using the WT-derived mitoEV protein list in the Reactome Pathway Browser[13]. Finally, an Enrichment Score was computed for each protein in the $App^{NL\text{-}G\text{-}F}$- versus WT-derived mitoEVs comparison by multiplying the -log10 (p-value) by the log2 fold change from the hybrid analysis. Genes were then ranked by Enrichment Score to perform Gene Set Enrichment Analysis (GSEA) using the previously defined custom annotation databases. GSEA was performed using the *fgsea* package, with an annotation size filter from 3 to 30,000, 100,000 permutations, and a p-value correction by FDR. All GSEA results were plotted as dot plots.

**Seahorse extracellular flux analysis**

Mitochondrial respiration and glycolytic function were assessed using a Seahorse XF96 extracellular flux analyzer (Agilent), which allows real-time measurement of oxygen consumption rate (OCR) and extracellular acidification rate (ECAR) in live cells and isolated mitochondrial fractions. Cells or vesicles were plated in poly-D-lysine or poly(ethyleneimine) (1:15,000; Sigma–Aldrich, #03880) coated 96-well Seahorse XF96 microplates, respectively (Agilent). The day before the assay, the Seahorse sensor cartridge was hydrated overnight in sterile dd$H_2O$ at 37 °C in a humidified, $CO_2$-free incubator. On the day of the experiment, dd$H_2O$ was replaced with XF calibrant solution (Agilent), previously equilibrated overnight in a $CO_2$-free incubator, and the cartridge was incubated for 1 h prior to calibration. Data were acquired using Wave 2.6.1 software (Agilent).

OCR in astrocytes: To assess mitochondrial respiration in astrocytes, culture medium was replaced with unbuffered DMEM supplemented with 17.5 mM glucose, 0.5 mM sodium pyruvate, and 2.5 mM L-glutamine (pH 7.2–7.4). OCR was measured at baseline and following sequential injections of 1 µM oligomycin A (Sigma-Aldrich), 1 µM FCCP (Sigma-Aldrich), and 0.5 µM antimycin A (Sigma-Aldrich) plus 0.5 µM rotenone (Sigma-Aldrich). Basal respiration, maximal respiration, and ATP-linked respiration were calculated using the Seahorse XF Cell Mito Stress Test report generator. OCR values are expressed as pmol $O_2$/min and normalized to protein content.

ECAR in astrocytes: ECAR was measured in astrocytes to evaluate glycolytic activity. Cells were incubated in unbuffered DMEM supplemented with 2.5 mM L-glutamine. ECAR was recorded at baseline and after sequential addition of 15 mM glucose, 1 μM oligomycin A, and 50 mM 2-deoxy-D-glucose. Glycolysis, glycolytic capacity, and glycolytic reserve were calculated using the Seahorse XF Glycolytic Stress Test Summary Report and expressed as mpH/min/μg protein.

Fatty acid β-oxidation–dependent OCR in neurons: To assess fatty acid oxidation (FAO)–dependent mitochondrial respiration in primary neurons, culture medium was replaced with unbuffered DMEM supplemented with 0.5 mM glucose, 1 mM L-glutamine, and 0.5 mM L-carnitine (pH 7.4). At the start of the assay, 1 mM palmitate conjugated to 0.17 mM BSA or 0.17 mM BSA alone (control) was added. OCR was measured at baseline and following sequential injections of 1 μM oligomycin A, 1 μM FCCP, and 0.5 μM antimycin A plus 0.5 μM rotenone. FAO-related respiratory parameters were calculated by subtracting BSA:Palm OCR to BSA OCR. Data are expressed as pmol $O_2$/min, normalized to protein content.

OCR in mitoEVs: Briefly, 15 μg of mitoEV protein was resuspended in 30 μL mitochondria assay solution (MAS: 70 mM sucrose, 220 mM mannitol, 10 mM $KH_2PO_4$, 5 mM $MgCl_2$, 1 mM EGTA, 2 mM HEPES-KOH, pH 7.2) supplemented with 0.2% fatty acid–free BSA and 8% polyethylene glycol (PEG600). MitoEVs were plated on coated XF96 plates and centrifuged at 2,200 *xg* for 20 min at 4 °C. MAS was further supplemented with 0.1 mM ADP (#A2754, Sigma-Aldrich), 10 mM pyruvate (#58636, Sigma-Aldrich), 2 mM malate (#46940-U, Sigma-Aldrich), pH 7.4, and 4 μM FCCP to support uncoupled respiration driven by Complex I. OCR was measured at baseline and following sequential injections of 2 μM rotenone (Complex I inhibitor), 10 mM succinate (Complex II substrate), 4 μM antimycin A (Complex III inhibitor), and 10 mM ascorbate plus 0.1 mM TMPD to assess Complex IV activity, as previously described[14].

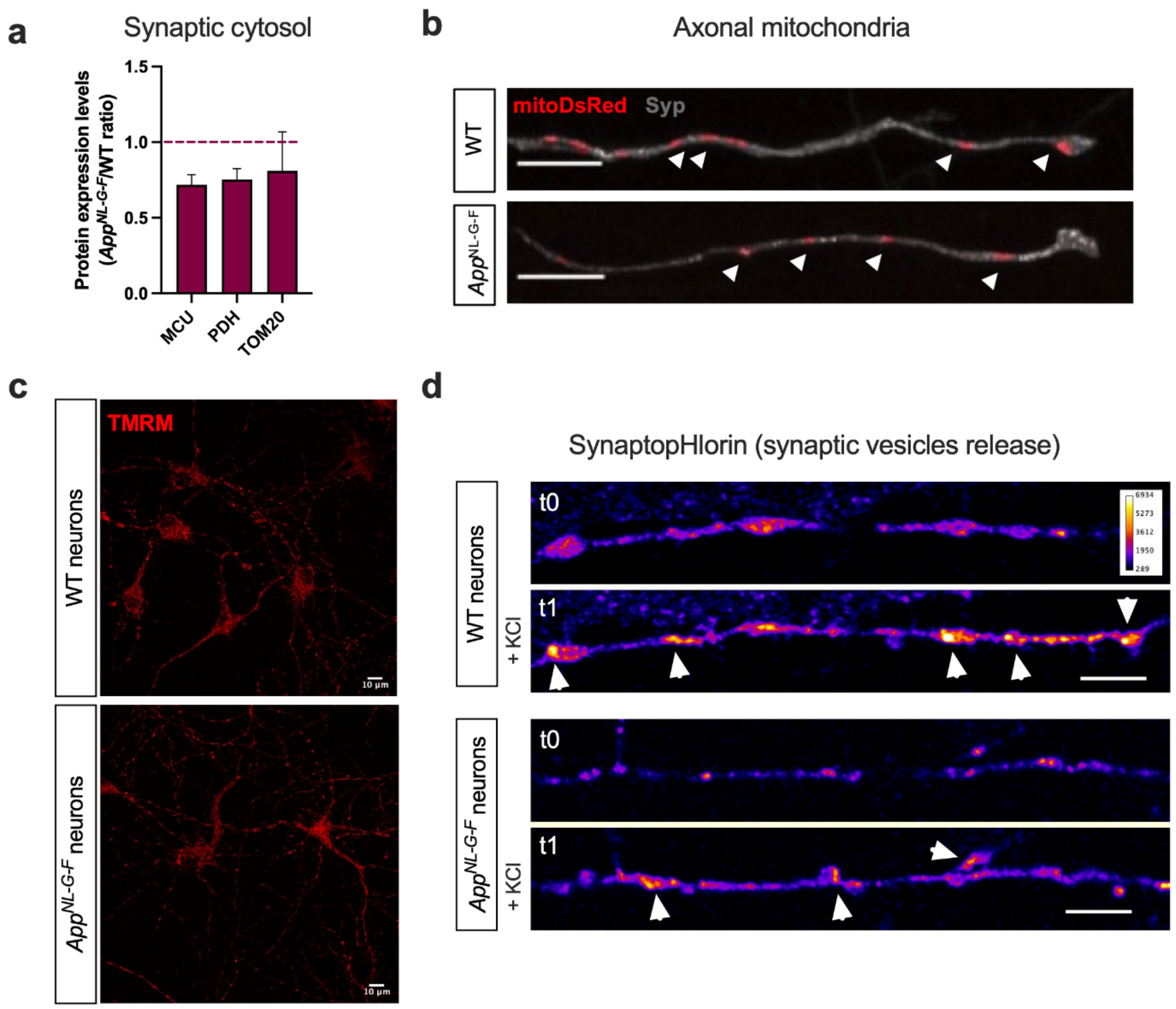


**Supplementary Fig S1: Characterization of *App*$^{NL-G-F}$ neuronal mitochondrial bioenergetics. (a)** *App*$^{NL-G-F}$/WT ratio of protein expression levels of the mitochondrial calcium uniporter (MCU), pyruvate dehydrogenase (PDH), and TOM20 in the presynapse (synaptic cytosol) (n = 3). **(b)** Representative confocal images of axonal mitochondria labelled with mitoDsRed and synaptophysin (Syp). Quantification is shown in Fig 1d. **(c)** Representative confocal images of mitochondrial membrane potential measured using TMRM under non-quenching conditions (10 nM, 30 min). Quantification is shown in Fig 1g. **(d)** Synaptic vesicle release was assessed in neurons expressing the synaptopHluorin reporter before and after stimulation with KCl (50 mM). Quantification is shown in Fig 1h, i. Scale bar = 10 µm.

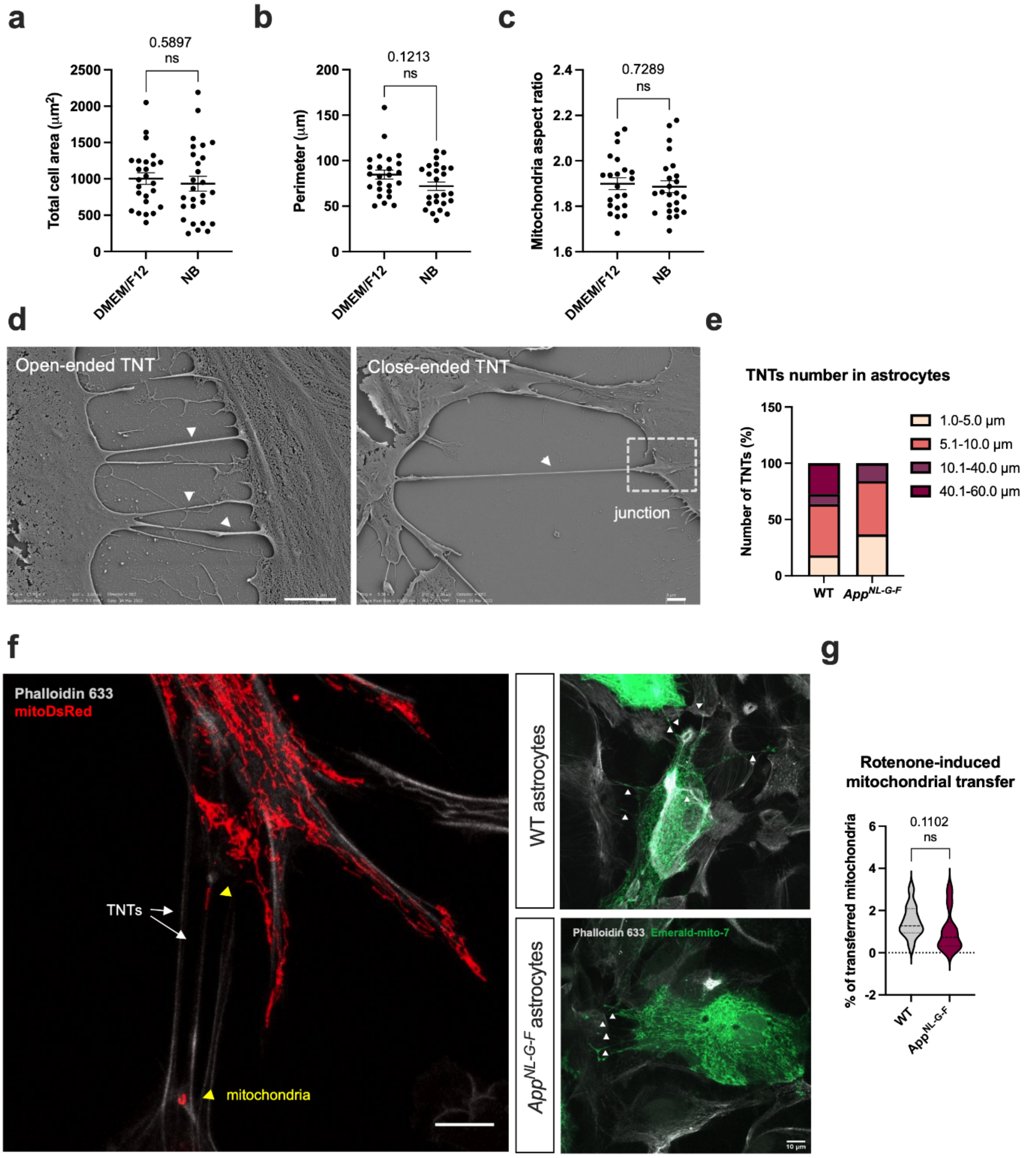


**Supplementary Fig S2: Effects of neurobasal media on astrocytes and intercellular mitochondria transfer via TNTs.** Astrocytic area **(a)** and perimeter **(b)**, and mitochondrial elongation **(c)** were quantified in astrocytes cultured in neuronal medium (Neurobasal, NB) or DMEM/F12 to assess potential effects of media change in co-culture conditions. **(d)** Representative scanning electron microscopy images depicting TNTs (arrows) between astrocytes. Left: Open-ended TNTs; Right: close-ended TNTs with a junction connected to the cell body. Scale bar = 3 µm. **(e)** Quantification of intercellular TNT length obtained from electron microscopy images. **(f)** A set of astrocytes were transfected with mitoDsRed or mitoEmerald and co-cultured with non-transfected astrocytes for 24 h. Actin filaments (including in TNT) were stained with phalloidin. Scale bar = 10 µm. **(g)** Before co-culture, non-transfected astrocytes were exposed to rotenone (50 nM, 24 h). Percentage of mitochondrial transfer to rotenone-treated astrocytes was quantified. ns. non-significant.

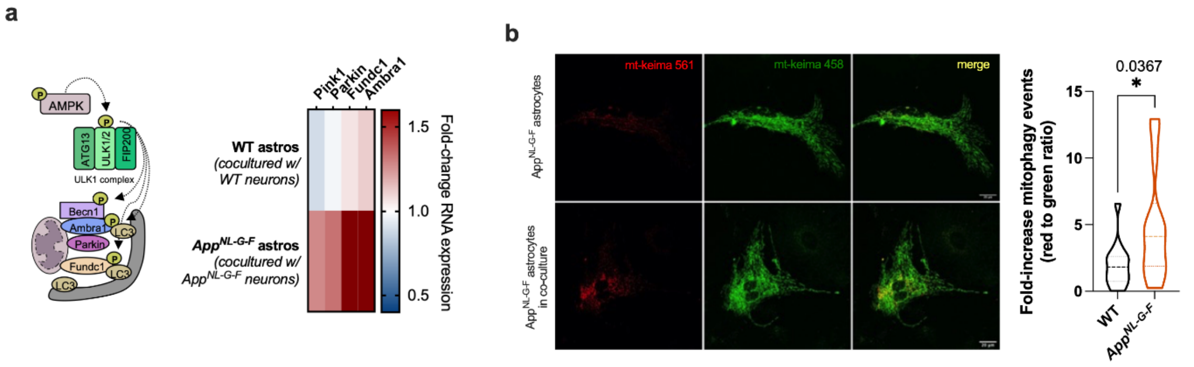


**Supplementary Fig S3: Upregulation of mitophagy events in *App*$^{NL\text{-}G\text{-}F}$ astrocytes in co-culture with neurons. (a)** Schematic representation of proteins involved in mitophagy regulation. Heatmap presents the expression levels of mitophagy-related genes in astrocytes in monoculture or co-culture (in Boyden chambers) with neurons. **(b)** Mt-Keima-transfected astrocytes in monoculture and co-culture with neurons. Data show fold-increase in mitophagy events in co-cultured astrocytes in comparison with monocultures. Statistical significance was analyzed using unpaired Mann-Whitney test; * $p \leq 0.05$.

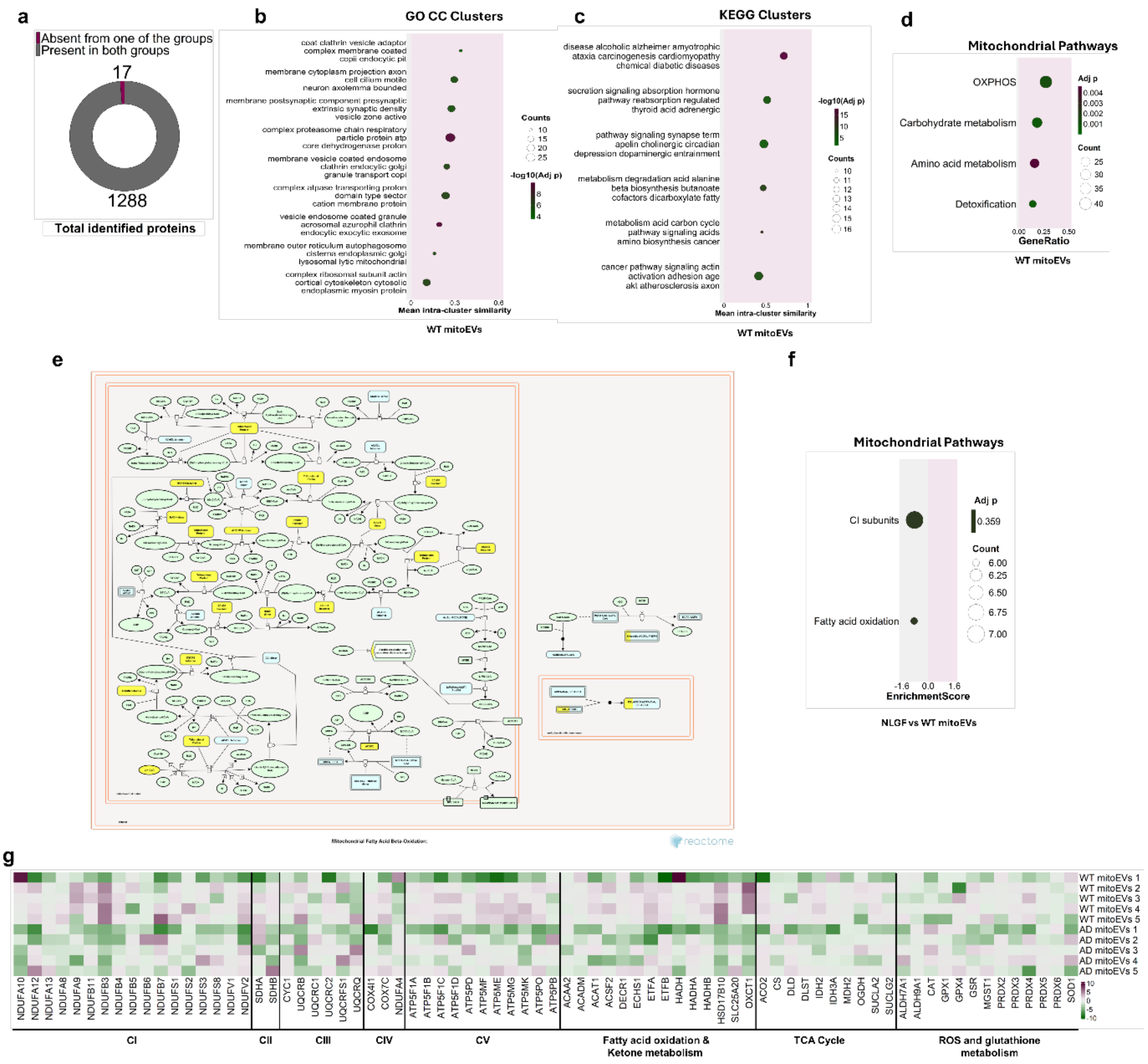


**Supplementary Figure S4: Comparative proteomic profiles and mitochondrial pathway enrichment in WT and *App*$^{NL\text{-}G\text{-}F}$ mitoEVs. (a)** Doughnut chart illustrating the number of identified proteins in WT and *App*$^{NL\text{-}G\text{-}F}$ astrocytic mitoEVs. Percentage of proteins detected in both genotypes (gray) versus percentage of proteins detected only in one of the groups (magenta). **(b-c)** ORA dotplots of **(b)** over-represented GO CC term clusters, **(c)** KEGG clusters and **(d)** mitochondrial pathways in WT astrocytic mitoEVs. **(e)** Reactome pathway map of mitochondrial FAO overlaid with proteins identified in the WT astrocytic mitoEV proteome. Yellow highlights detected proteins. **(f)** Dotplot of GSEA depicting the *App*$^{NL\text{-}G\text{-}F}$ versus WT astrocytic mitoEV comparison, focusing on mitochondrial pathways subterms. Dot size and color represent the number of proteins associated with each term and the adjusted p-value, respectively (applies to all dot plots). **(g)** Heatmap of the mitochondrial pathway subterm-related proteins in WT or *App*$^{NL\text{-}G\text{-}F}$ astrocytic mitoEVs. The columns represent the proteins and the rows the samples. The color gradient encodes the column-centered intensities of each protein across all samples.

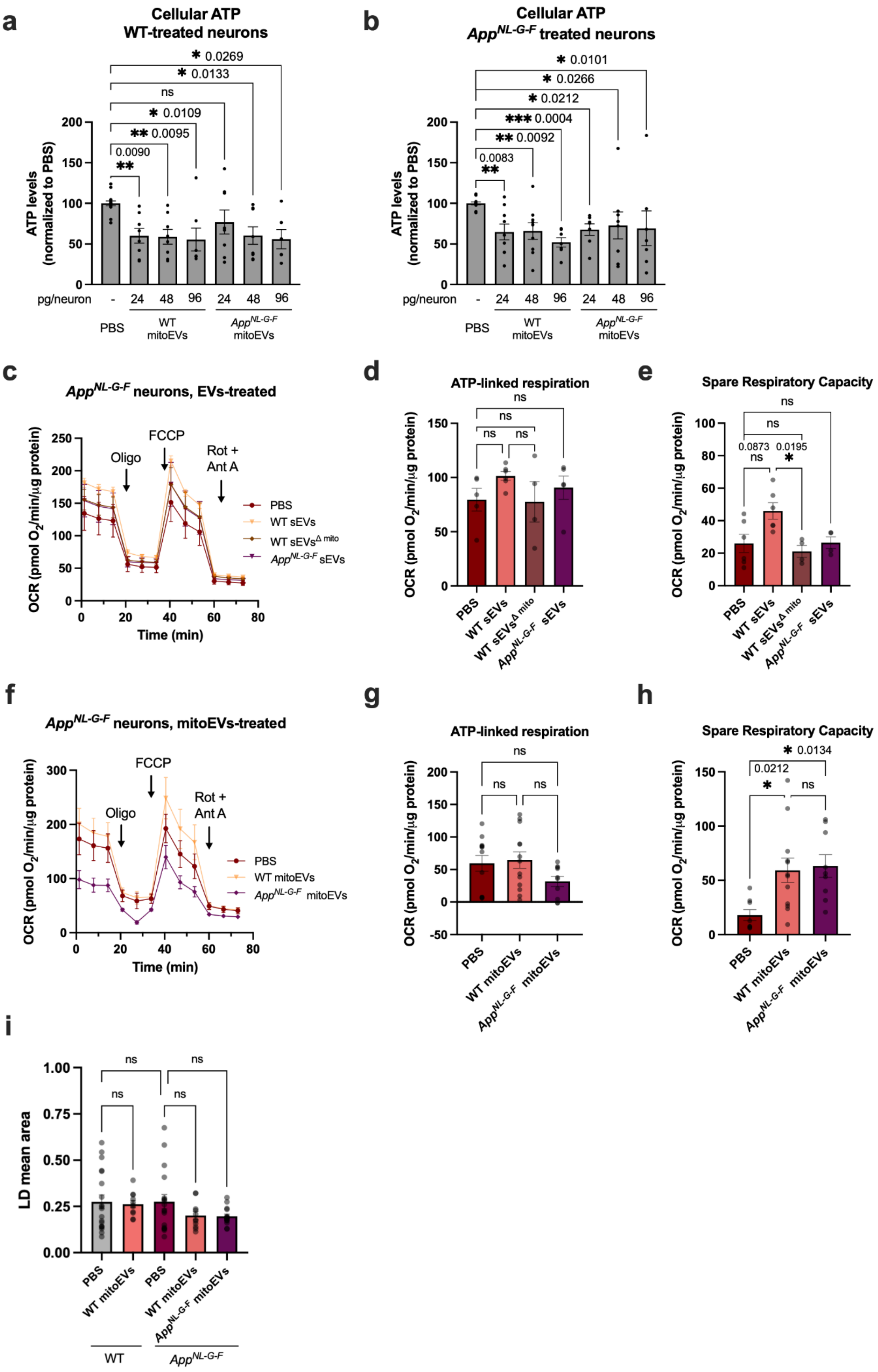

a
Cellular ATP
WT-treated neurons
* 0.0269
* 0.0133
ns
* 0.0109
** 0.0095
0.0090
**
ATP levels (normalized to PBS)
pg/neuron
-
24
48
96
PBS
WT mitoEVs
App^NL-G-F mitoEVs
b
Cellular ATP
App^NL-G-F treated neurons
* 0.0101
* 0.0266
* 0.0212
*** 0.0004
** 0.0092
0.0083
**
c
App^NL-G-F neurons, EVs-treated
OCR (pmol O2/min/µg protein)
Oligo
FCCP
Rot + Ant A
PBS
WT sEVs
WT sEVs^Δ mito
App^NL-G-F sEVs
Time (min)
d
ATP-linked respiration
ns
e
Spare Respiratory Capacity
0.0873
0.0195
f
App^NL-G-F neurons, mitoEVs-treated
WT mitoEVs
App^NL-G-F mitoEVs
g
h
* 0.0134
0.0212
i
LD mean area
WT
App^NL-G-F

**Supplementary Figure S5: MitoEVs effects on cellular ATP levels and respiration.** Neurons were treated, when indicated, with astrocytes-derived EVs (0.5 ng/neuron, 6 h) or mitoEVs (24, 48 or 96 pg/neuron, 24 h) (n = 5-8). **(a, b)** Cellular ATP levels were quantified using CellTiter-Glo in (a) WT and (b) $App^{NL\text{-}G\text{-}F}$ treated neurons. **(c-h)** OCR in EV- (c-e) and mitoEV (f-h)-treated neurons was evaluated using the Seahorse apparatus (n = 5-7). Functional mitoEVs were depleted from the EV pool ($EVs^{\Delta mito}$) by treating the EVs with the mitochondria uncoupler FCCP (2 μM, 30 min). Data presented as mean ± SEM, and statistical significance was done using Kruskal-Wallis test. * p ≤ 0.05, ** p ≤ 0.01, *** p ≤ 0.001, ns. non-significant.

**Supplementary Table S1:** List of qPCR primers from Applied Biosystems

| **Primers** | **Sequence code** |
|---|---|
| **Atp5k** | Mm00833200_g1 |
| **mt-Atp8** | Mm04225236_g1 |
| **Kif5b** | Mm00515276_m1 |
| **Ppargc1a** | Mm01208835_m1 |
| **Rhot1** | Mm01304158_m1 |
| **Rhot2** | Mm00524478_m1 |
| **Tfam** | Mm00447485_m1 |
| **Ambra1** | Mm00554370_m1 |
| **Fundc1** | Mm00511132_m1 |
| **Park2** | Mm00450186_m1 |
| **Pink1** | Mm00550827_m1 |
| **Fis1** | Mm00481580_m1 |
| **Dmn1l** | Mm01342903_m1 |
| **Mff** | Mm01273401_m1 |
| **Mtfr1** | Mm01289320_m1 |
| **Mtfr2** | Mm00510872_m1 |
| **Mief1** | Mm00724569_m1 |
| **Mief2** | Mm01234249_g1 |
| **Tubb3** | Mm00727586_s1 |

**Supplementary Table S2:** List of qPCR primers from Applied Biosystems

| Antibody | Dilution | Brand | Cat# |
|---|---|---|---|
| | **Primary antibodies** | | |
| **PSD95** | 1:1000 | Abcam | ab2723 |
| **Synaptophysin** | 1:1000 | Abcam | ab14692 |
| **GAPDH** | 1:1000 | Abcam | ab8245 |
| **OXPHOS** | 1:1000 | Abcam | ab110413 |
| **TOM20** | 1:1000 | Santa Cruz Biotechnology | sc-11415 |
| **MCU** | 1:1000 | Sigma Aldrich | HPA016480 |
| **PDH-E1α** | 1:1000 | Santa Cruz Biotechnology | sc-377092 |
| **OPA1** | 1:1000 | BD Bioscience | 612606 |
| **Alix** | 1:500 | Cell Signaling Technology | 2171S |
| **Flotillin-1** | 1:500 | BD Biosciences | 610820 |
| **ITM2B/Bri2** | 1:200 | Santa Cruz Biotechnology | sc-374362 |
| **FAM49B** | 1:200 | Proteintech | 20127-1-AP |
| **HSD17B10/ERAB** | 1:500 | Thermo- Fisher | MA5-42639 |
| **Mrps25** | 1:200 | Proteintech | 15277-1-AP |
| **VDAC1/2/3** | 1:1000 | Abcam | ab15895 |
| **β3-tubulin** | 1:5000 | Abcam | ab6046 |
| | **Secondary antibodies** | | |
| **IRDye 800CW anti-mouse** | 1:20,000 | Invitrogen | 926-32212 |
| **IRDye 800CW anti-rabbit** | 1:20,000 | Invitrogen | 926-32213 |
| **IRDye 680CW anti-mouse** | 1:20,000 | Invitrogen | |
| **IRDye 680CW anti-rabbit** | 1:20,000 | Invitrogen | |
| **Alexa Fluor 594 anti-rabbit** | 1:500 | Invitrogen | A11012 |

**Supplementary Table S3:** List of reagents and kits used

| Material | Manufacturer | Category number |
|---|---|---|
| **Isoflurane** | Abbott Scandinavia AB | 5260 |
| **Hibernate-E medium** | Thermo Fisher | A1247601 |
| **Hank's Balanced Salt Solution** | Thermo Fisher | 14175095 |
| **Trypsin/EDTA** | Thermo Fisher | 25200072 |
| **Fetal Bovine Serum** | Thermo Fisher | A5256801 |
| **DNAse I** | Roche | 10104159001 |
| **B-27 Supplement (50X), serum free** | Thermo Fisher | 17504044 |
| **GlutaMAX Supplement** | Thermo Fisher | 35050061 |
| **DMEM/F12 Cell Culture Media** | Thermo Fisher | 31331093 |
| **Poly-D-Lysine** | Sigma Aldrich | P6407 |
| **N2 supplement** | Thermo Fisher | 17502001 |
| **PLX-3397** | MedChemExpress | HY-16749 |
| **Lipofectamine 2000** | Thermo Fisher | 11668019 |
| **CellTracker Blue** | Thermo Fisher | C12881 |
| **Tetramethylrhodamine, Methyl Ester, Perchlorate (TMRM)** | Thermo Fisher | T668 |
| **CellTiter-Glo® Luminescent Cell Viability Assay** | Promega | G7570 |
| **DMEM 5030 media** | Sigma-Aldrich | 5030 |
| **Sodium pyruvate (100 nM)** | Thermo Fisher | 11360070 |
| **RNeasy® Mini Kit** | Qiagen | 74104 |
| **High-Capacity cDNA Reverse Transcription Kit** | Applied Biosystems | 4368814 |
| **TaqMan™ Fast Advanced Master Mix** | Applied Biosystems | 4444557 |
| **Etomoxir** | Sigma-Aldrich | 236020 |
| **DPBS, no calcium, no magnesium** | Thermo Fisher | 14190250 |
| **OptiPrep™** | StemCell Technologies | 07820 |
| **Carbonyl cyanide-4-(trifluoromethoxy) phenylhydrazone (FCCP)** | Sigma-Aldrich | C2920 |
| **NuPAGE MES SDS running buffer** | Invitrogen | NP0002 |
| **EZBlue™ Gel Staining Reagent** | Sigma-Aldrich | G1041 |
| **Benzonase® Nuclease** | Millipore | 70664-10KUN |
| **Pierce BCA Protein Assay Kit** | Thermo Fisher | 23225 |
| **ECL™ Rainbow™ Marker – Full Range** | GE HealthCare | RPN800E |
| **Ponceau staining** | Sigma-Aldrich | P7170 |
| **Triton X-100** | Sigma-Aldrich | X100-1L |
| **Bovine Serum Albumin** | Sigma-Aldrich | A6003 |
| **Phalloidin-633 probe** | Thermo Fisher | A22284 |
| **Antifade mounting media** | VectaShield Plus | H-1900 |
| **Poly(ethyleneimine)** | Sigma-Aldrich | 03880 |

| | | |
|---|---|---|
| **XF calibrant solution** | Agilent | 100840-000 |
| **Oligomycin A** | Sigma-Aldrich | 75351 |
| **Antimycin A** | Sigma-Aldrich | A8674 |
| **Rotenone** | Sigma-Aldrich | R8875 |
| **Adenosine 5′-diphosphate sodium salt (ADP)** | Sigma-Aldrich | A2754 |
| **Sodium pyruvate solution** | Sigma-Aldrich | 58636 |
| **D-Malic acid** | Sigma-Aldrich | 46940-U |
| **BODIPY™ 493/503** | Thermo Fisher | D3922 |
| **CellTracker™ Red CMTPX** | Thermo Fisher | C34552 |
| **Neurobasal Medium, phenol red–free** | Thermo Fisher | 12348017 |

**Supplementary Table S4:** List of materials used

| Material | Manufacturer | Category number |
|---|---|---|
| **Microfluidic chambers** | Xona Microfluidics | DOC450 |
| **Boyden chambers** | Falcon Corning Life Sciences | 353095 |
| **mPES 300kD hollow fiber membrane** | Repligen | D06-E300-05-N |
| **38.5 mL, Open-Top Thinwall Ultra-Clear Tube** | Beckman Coulter | 344058 |
| **14 mL Open-Top Thinwall Ultra-Clear Tube** | Beckman Coulter | 344060 |
| **Bis-Tris gel 12%** | Invitrogen | NP0341BOX |
| **NuPAGE 4-12 % Bis-Tris gels** | Invitrogen | NP0321BOX |
| **Amersham™ Protran® Western blotting nitrocellulose membrane** | GE HealthCare | 10600002 |
